# Peri-mitochondrial actin filaments inhibit Parkin assembly via disruption of ER- mitochondrial contact

**DOI:** 10.1101/2024.07.26.605389

**Authors:** Tak Shun Fung, Amrapali Ghosh, Marco Tigano, Henry N Higgs, Rajarshi Chakrabarti

## Abstract

Mitochondrial damage represents a dramatic change in cellular homeostasis, necessitating metabolic adaptation as well as clearance of the damaged organelle. One rapid response to mitochondrial damage is peri-mitochondrial actin polymerization within 2 mins, which we term ADA (<u>A</u>cute <u>D</u>amaged-induced <u>A</u>ctin). ADA is vital for a metabolic shift from oxidative phosphorylation to glycolysis upon mitochondrial dysfunction. In the current study we investigated the effect of ADA on Pink1/Parkin mediated mitochondrial quality control. We show that inhibition of proteins involved in the ADA pathway significantly accelerates Parkin recruitment onto depolarized mitochondria. Addressing the mechanism by which ADA resists Parkin recruitment onto depolarized mitochondria, we found that ADA disrupts ER-mitochondrial contacts in an Arp2/3 complex-dependent manner. Interestingly, over-expression of ER-mitochondrial tethers overrides the effect of ADA, allowing rapid recruitment of not only Parkin but also LC3 after mitochondrial depolarization. During chronic mitochondrial dysfunction, Parkin and LC3 recruitment are completely blocked, which is reversed rapidly by inhibiting ADA. Taken together we show that ADA acts as a protective mechanism, delaying mitophagy following acute damage, and blocking mitophagy during chronic mitochondrial damage.

## Introduction

Mitochondria are well known to oxidize organic molecules for ATP production, using the mitochondrial membrane potential (Δψm) generated by the electron transport chain (ETC) across the inner mitochondrial membrane (IMM) to drive ATP synthase (Mitchell, 2011). In addition, mitochondria are important signaling and biosynthetic hubs, participating in calcium signaling (Nicholls, 2005), lipid synthesis (Nowinski *et al*, 2020), amino acid synthesis(Ahn & Metallo, 2015), iron-sulfur cluster biosynthesis (Braymer & Lill, 2017), and apoptosis (Green, 2022). Mitochondrial dysfunction can lead to unregulated cell death *via* ferroptosis (Gao *et al*, 2019) as well as activation of host-pathogen responses and innate immunity (Kim *et al*, 2023). For these reasons, several pathways exist to mitigate mitochondrial dysfunction, including mitochondrial destruction by mitophagy.

Mammalian mitophagy involves recognition of the dysfunctional mitochondrion through specific receptors and/or adaptors, often in a trigger-specific manner. One such trigger is depolarization (the sustained loss of Δψm), a common outcome of mitochondrial dysfunction. Mitochondrial depolarization activates the Pink1/Parkin mitophagy pathway, which in turn stimulates recruitment of LC3 to feed the mitochondrion into the autophagy pathway. In a healthy mitochondrion, the serine-threonine protein kinase Pink1 is continuously degraded through a system that involves its import into the mitochondrial matrix through the Tom and Tim complexes, which requires mitochondrial membrane potential (Jin *et al*, 2010) (Deas *et al*, 2011) (Meissner *et al*, 2011)(Yamano & Youle, 2013). Upon mitochondrial depolarization, Pink1 is stabilized onto the OMM, where it homodimerizes, self-activates, and phosphorylates both ubiquitin and Parkin (a cytoplasmic E3 ubiquitin ligase), leading to Parkin accumulation on OMM (Lazarou *et al*, 2012; Okatsu *et al*, 2013). Mitochondrially-recruited Parkin ubiquitinates several substrates, generating an ubiquitinated mask on the OMM which is recognized by a group of adaptors (P62, NBR1, NDP52, TAX1BP1 and Optineurin (OPTN)). These adaptor proteins subsequently recruit LC3.

A growing number of studies have documented several important roles for actin filaments in regulation of mitochondrial homeostasis and dynamics (Li *et al*, 2015; Moore *et al*, 2021, 2021; Basu *et al*, 2021; Fung *et al*, 2023; Chakrabarti *et al*, 2018; Korobova *et al*, 2013). We and others have shown that acute mitochondrial depolarization using uncouplers like FCCP or CCCP generates a transient cloud of actin filaments, which we term ADA (<u>a</u>cute <u>d</u>amage-induced <u>a</u>ctin), around the depolarized mitochondrion within 5 min (Li *et al*, 2015; Fung *et al*, 2019) through signaling pathways requiring AMP-dependent protein kinase (AMPK) and protein kinase C (Fung *et al*, 2022a). ADA is also activated by hypoxia, ETC inhibitors, and ATP synthase inhibition (Chakrabarti *et al*, 2022). Interestingly, chronic mitochondrial dysfunction (through depletion of mtDNA or the key ETC protein NDUFS4) causes ADA-like filaments around mitochondria that persist for days but are constantly turning over, with a half-life of less than 10 min (Chakrabarti *et al*, 2022).

Two functional consequences for ADA have been identified. ADA transiently suppresses mitochondrial morphological dynamics that are triggered by mitochondrial depolarization (Fung *et al*, 2019, 2022a). These dynamics involve rearrangement of the inner mitochondrial membrane (IMM), dependent on the IMM protease Oma1, resulting in circularization of the IMM within an intact outer mitochondrial membrane (OMM) (Fung *et al*, 2022a, 2019; Miyazono *et al*, 2018). ADA inhibits both mitochondrial circularization and proteolytic processing of the Oma1 substrate Opa1. A second ADA function is to rapidly stimulate glycolysis. This glycolytic stimulus also is a feature of the persistent ADA-like filaments that assemble following chronic mitochondrial damage (Chakrabarti *et al*, 2022). It is possible that ADA has additional effects.

Here, we asked whether ADA has a role in regulating mitophagy. We show that ADA significantly delays Parkin recruitment and downstream LC3 recruitment, through acutely disrupting endoplasmic reticulum (ER)-mitochondrial contacts (ERMC). Intriguingly, disassembly of the ADA-like filaments present around chronically damaged mitochondria allows both Parkin and LC3 recruitment. Overall, our results suggest that ADA serves as an acute roadblock to the early events in Pink1-Parkin mediated mitophagy, through the regulation of ER-mitochondrial contacts.

## Results

### ADA delays Parkin recruitment onto damaged mitochondria

Damaged or depolarized mitochondria can be subject to multiple fates, including mitophagy (Boland *et al*, 2013; Palikaras *et al*, 2018). Though the final steps of mitophagy occur much later than the initial depolarization (McWilliams & Muqit, 2017), Parkin recruitment to the OMM is an early step in the process (Harper *et al*, 2018; Pickles *et al*, 2018).To test whether induction of ADA affected recruitment of Parkin to damaged mitochondria, we blocked ADA by inhibiting the Arp2/3 complex and evaluated the kinetics of GFP-Parkin recruitment to mitochondria by live-cell microscopy and compared it with that of control cells. In control U2-OS cells, Parkin recruitment starts at 45.5 min ± 10.7 min after CCCP treatment. Both CK666 pre-treatment (Arp2/3 inhibition) (**Fig 1A-C**) and Arp2 KD (Arp2/3 depletion) (**Fig 1D-F; *EV1A (upper)***) result in significant acceleration of Parkin recruitment (29.4 min ± 9.0 min and 31.6 min ± 8.3 min, respectively). We obtain similar results in HeLa cells, where CK666 treatment (***Fig EV1B- D***) or Arp2 KD (***Fig EV1A (lower);***) causes significantly faster CCCP-induced Parkin recruitment (25.2 min ± 10.9 min and 29.0 min ± 6.1 min, respectively) than control cells (46.3 min ± 12.2 min). To test this effect further, we suppressed two other key components of the ADA signaling pathway: the Wave Regulatory Complex (WRC) and the FMNL formins (Fung *et al*, 2022b). Either FMNL KD (32.6 min ± 8.7 min) or WRC KD (33.4 min ± 10.3 min) in U2OS cells causes significant acceleration of Parkin recruitment compared to control cells (**Fig 1D-F; *EV1A (upper)***). FMNL depletion in HeLa cells also showed a similar trend of accelerated Parkin recruitment compared to control cells (**Fig *EV1E-G; Fig EV1A (lower)***). We also used a fixed-cell assay to evaluate the number of HeLa cells with GFP-Parkin positive mitochondria after treatment with CCCP for 30 minutes. In accordance with live cells data, we detected a significantly higher proportion of mitochondrial GFP-positive cells in Arp2 KD (72.2 ±25.5 %) compared to control (32.3 ±22.3 %) (***Fig EV1H, I)***. Taken together, these results suggest that ADA significantly delays Parkin recruitment onto damaged mitochondria in both U2-OS and HeLa cells.

**Figure 1:**
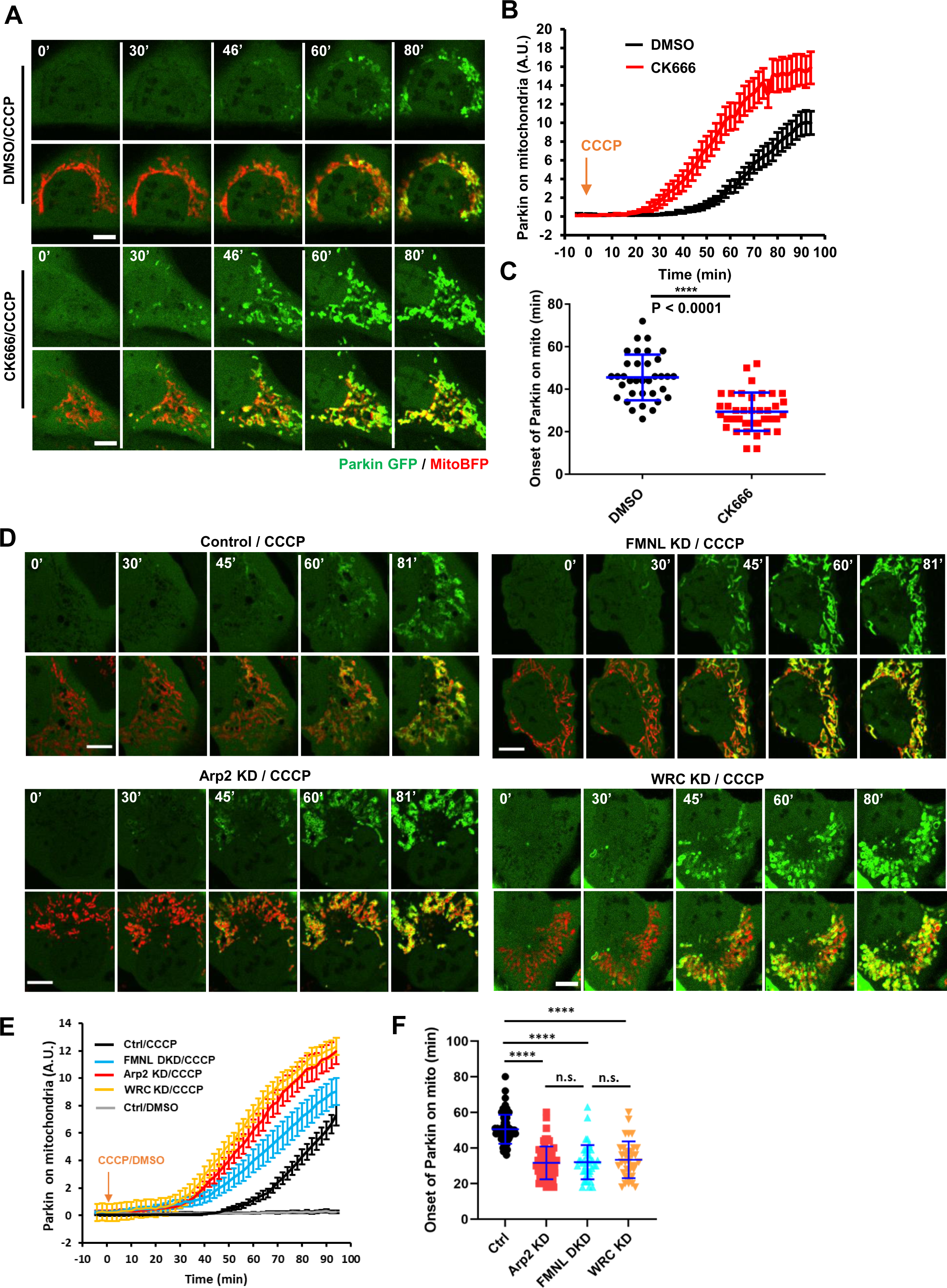
ADA delays Parkin recruitment to depolarized mitochondria, (A) Time lapse montages of WT U2OS cells transfected with GFP-Parkin (green) and Mito-BFP (red) pre-treated with 30 mins 100μM CK666 or DMSO before treatment with 20μM CCCP treatment during live-cell imaging at time 0. Imaging conducted at the medial cell section. Scale: 10μm. Time in mins. (**B**) Quantification of colocalized Parkin signal with mitochondrial signal in live-cell imaging after 20μM treatment of CCCP at time 0 for 30 mins DMSO (black) or 100μM CK666 (red) pre-treated U2OS cells. N = 35 cells for DMSO/CCCP and 38 for CK666/CCCP. 4 independent experiments. Error ± SEM **(C)** Scatter plot of Parkin signal onset on mitochondria for DMSO or 100μM CK666 treated U2OS cells. Parkin onset time = 45.54 min ± 10.74 (mean ± s.d.) for DMSO/CCCP and 29.42 ± 10.74 for CK666/CCCP. Error: ±SD. Same dataset as in Figure 1B. P< 0.0001 (****). Student’s unpaired t-test. **(D)** Time lapse montages of ctrl, Arp2 KD, FMNL1/3 DKD and WRC KD U2OS cells transfected with GFP-Parkin (green) and Mito- BFP (red) treated with 20μM CCCP treatment at time 0 during live-cell imaging. Imaging conducted at the medial cell section. Scale: 10μm. Time in mins. (**E**) Quantification of colocalized Parkin signal with mitochondrial signal in live-cell imaging after 20μM treatment of CCCP at time 0 for ctrl, FMNL1/3 DKD or Arp2 KD U2OS cells. N = 79 cells for ctrl; 62 for Arp2 KD; 42 for FMNL1/3 DKD and 35 for WRC KD. 3 independent experiments. Error ± SEM **(F)** Scatter plot of Parkin signal onset on mitochondria for ctrl, Arp2 KD, FMNL1/3 DKD and WRC KD in U2OS cells. Parkin onset time = 46.97 min ± 8.86 (mean ± s.d.) for ctrl; 32.06 ± 9.03 for Arp2 KD; 32.61 ± 8.75 for FMNL 1/3 DKD and 33.37 ± 10.30 for WRC KD cells. Same dataset as in Figure 1E. P< 0.0001 (****) for ctrl *vs* FMNL1/3 DKD and ctrl *vs* Arp2 KD; P = 0.9795 (n.s.) for FMNL1/3 DKD *vs* Arp2 KD, P = 0.7808 (n.s.) for WRC KD *vs* Arp2 KD. Tukey’s multiple comparisons test used. Error: ± SD

Next, we investigated the mechanism by which ADA delays Parkin recruitment. One possibility is that the thick network of actin filaments serves as a diffusive barrier to impede access of Parkin or its upstream factor(s) to the OMM. To test this possibility, we utilized an inducible mitochondrial recruitment system, by expressing cytosolic CFP-(FRB)5 and mitochondrially targeted AKAP1-FKBP-YFP. Upon rapamycin addition, CFP-(FRB)5 is recruited to the OMM in 15-30 sec (***Fig EV2A***). We tested the effect of ADA on CFP-(FRB)5 recruitment by adding rapamycin ∼100 sec after CCCP addition (the point of maximal ADA (Fung *et al*, 2019, 2022b). The rate of CFP-(FRB)5 recruitment is not measurably influenced by ADA (***Fig EV2B, C***). Since CFP-(FRB)5 is larger than Parkin (82 *versus* 52 kDa), these results suggest that the actin filaments polymerized during ADA do not physically block the access of Parkin to depolarized mitochondria. We therefore examined other mechanisms that could explain the delay in Parkin recruitment.

### ADA disrupts ER-mitochondrial contacts

Curiously, we noticed that the actin filaments assembled during ADA often occurred between mitochondria and ER (***Fig EV2D***). We therefore asked whether ADA could actively displace ER or other organelles from depolarized mitochondria. By imaging simultaneously mitochondria, ER and actin filaments in live cells, we recorded a significant reduction in the overlap between ER and mitochondria (ERMCs) occurring with a similar kinetic to that of CCCP induced ADA (**Fig 2A, B**). Importantly, the acute decrease in ER-mitochondrial overlap depends on actin assembly, as treatment with CK666 restores ER-mitochondrial overlap upon CCCP induced depolarization (**Fig 2B**). As a control, we tested whether ADA could disrupt the contact between mitochondria and another subcellular organelle, lysosomes. The extent of mitochondria and lysosome contacts was not influenced by ADA (***Fig EV2E, F***). Altogether, these results suggest that ADA might specifically disrupt ERMCs.

**Figure 2:**
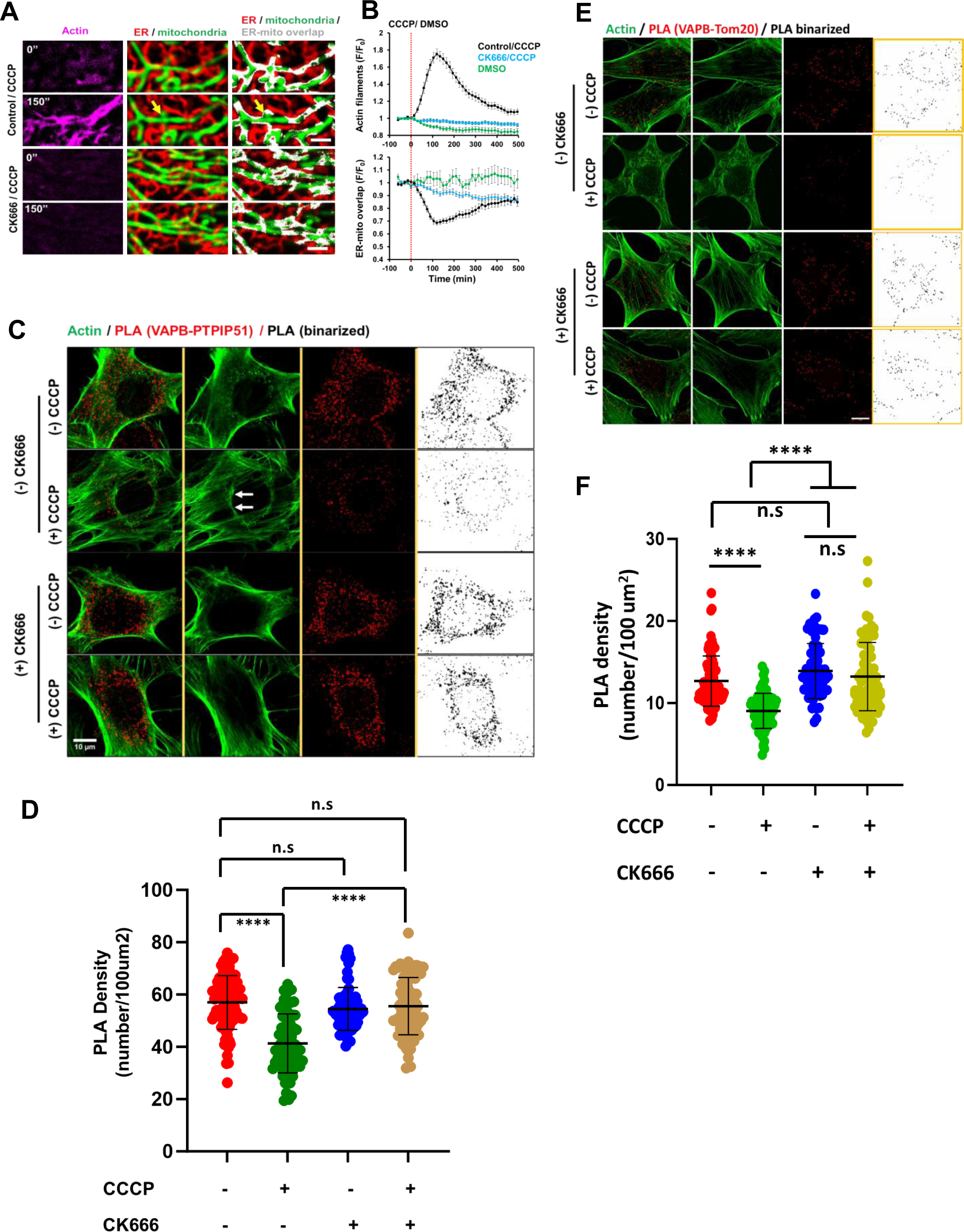
ADA acutely disrupts ER-Mitochondrial contacts (ERMC), (A) Time lapse montages of CCCP-induced actin polymerization and ER-mitochondrial overlap (grey) in U2OS cells transfected with ER-RFP (ER; red), mito-BFP (mitochondria; green) and GFP-F tractin (actin filaments; magenta) and pre-treated with either DMSO or CK666 (100 μM; 30 min). Scale: 10 µm. Time in mins (B) Quantification of CCCP-induced actin polymerization and ER-mitochondrial overlap in U2OS cells transfected with ER-RFP, mito-BFP and GFP-F tractin and pre-treated with either DMSO or CK666 (100 μM; 30 min). N= 45 cells (CCCP), 34 cells (CK666/CCCP) and 19 cells (DMSO). 3 independent experiments. Error ± SEM (C) Representative fixed-cell images from Proximity Ligation Assay (PLA) conducted between VAPB (ER) and PTPIP51 (mitochondria) to assess ERMC (red and binarized) in HeLa cells treated with 20 μM CCCP for 5 min in the presence or absence of CK666 (100 μM). Cells were stained with Phalloidin-488 to label actin filaments prior to PLA staining. Arrows represent ADA induction. Scale: 10 µm. (D) Scatter plot representing density of PLA dots (no. per 100µm^2^) in control and CCCP-treated HeLa cells in the presence or absence of CK666. Same data set as Fig 2C. N= 86 cells (control); 72 cells (CCCP); 75 cells (CK666) and 84 cells (CK666 + CCCP) from 20-25 fields and 3 independent experiments. P< 0.0001 (****) for untreated vs CCCP and CCCP vs CK666/CCCP. Error: ±SD (E) Representative fixed-cell images from Proximity Ligation Assay (PLA) conducted between VAPB (ER) and Tom20 (mitochondria) to assess ERMC (red and binarized) in HeLa cells treated with 20 μM CCCP for 5 min in the presence or absence of CK666 (100 μM). Cells were stained with Phalloidin-488 to label actin filaments prior to PLA staining. Arrows represent ADA induction. Scale: 10 µm. (F) Scatter plot representing density of PLA dots (no. per 100µm^2^) in control and CCCP-treated HeLa cells in the presence or absence of CK666. Same data set as Fig 2E. N= 80 cells (control); 85 cells (CCCP); 65 cells (CK666) and 77 cells (CK666 + CCCP) from 20-25 fields and 2 independent experiments. P< 0.0001 (****) for untreated vs CCCP and CCCP vs CK666/CCCP. Error: ±SD

To test the effect of ADA on ERMCs in a more direct manner, we used Proximity Ligation Assay (PLA) for interacting partners that mediate ERMCs: VAP-B (ER) and PTPIP51 (mitochondria)(Obara *et al*, 2024). Mouse Embryonal Fibroblasts (MEFs) treated for 5 min with CCCP displayed ADA induction, as well as a significant decrease in PLA density (**Fig 2C, D**) suggesting an acute reduction of ERMC. Consistently, simultaneous treatment of CK666 together with CCCP restores PLA density (**Fig 2C**), while acute CK666 treatment alone does not result in significant changes in PLA density compared to the control cells (**Fig 2D**). We repeated the PLA assay with a second combination of proteins: VAP-B (ER) and Tom20 (mitochondria). Again, CCCP causes a decrease in PLA density prevented by CK666 (**Fig 2E, F**). Overall, these results suggest that peri-mitochondrial actin assembly following acute mitochondrial depolarization dynamically disrupts ERMC.

### ADA-induced ERMC disruption delays Parkin recruitment onto mitochondria

In light of previous reports suggesting that a sub-set of Parkin recruitment occurs at ERMCs (Chen & Dorn, 2013; Gelmetti *et al*, 2017) we tested whether increasing ER-mitochondrial contacts could overcome the inhibitory effect of ADA on Parkin recruitment. To increase ERMCs, we over-expressed VAP-B (Gomez-Suaga *et al*, 2017). VAP-B overexpression (***Fig EV3A***) accelerates mitochondrial recruitment of Parkin upon CCCP treatment to an extent similar to that of Arp2- or FMNL-depletion (**Fig 3A-C**), without affecting the kinetics of ADA (***Fig EV3B***). To exclude the possibility of a specific effect of VAP-B on stimulating Parkin recruitment by direct interaction, we also tested a synthetic ER-mitochondrial linker (*ER-RFP-Mito*) (Nichtová *et al*, 2023). *ER-RFP-Mito* enhances histamine-induced ER-to-mitochondria calcium transfer (***Fig EV3C, 3D***), indicative of an increase in functional ERMCs. *ER-RFP-mito* also results in a significantly higher proportion of FCCP-treated cells displaying Parkin-positive mitochondria (49.7 ± 5.8% and 67.7 ± 5.9% in 15 and 30 minutes respectively) over control cells (13.5 ±3.6% and 32.5 ±8.6%), (**Fig 3D, E**). Notably, *ER-RFP-mito* expression does not inhibit CCCP-induced ADA (***Fig EV3E***).

**Figure 3:**
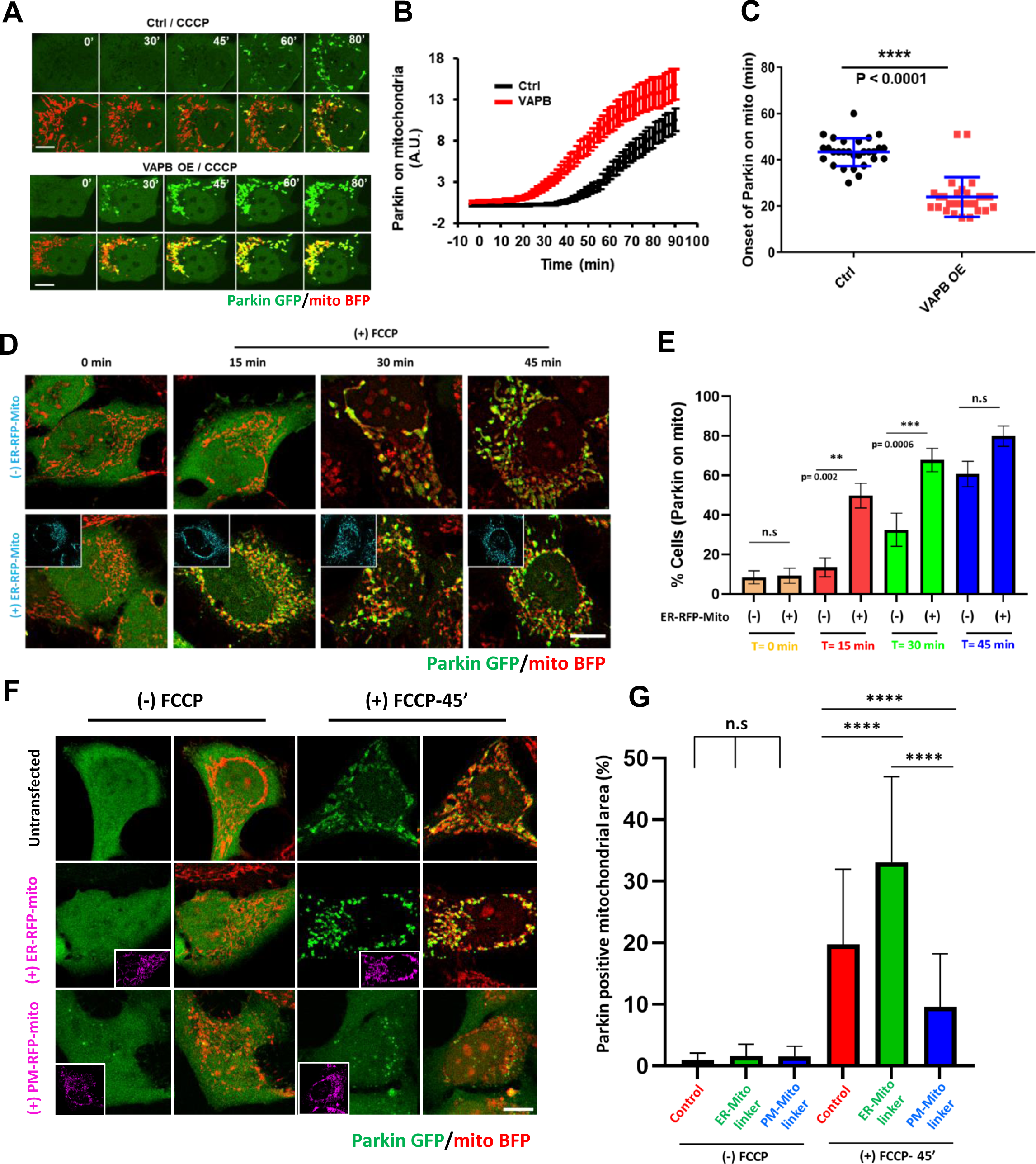
Enhancing ERMC speeds up Parkin recruitment on depolarized mitochondria. (A) Time lapse montages of control and myc-VAP-B-overexpressing U2OS cells transfected with GFP-Parkin (green) and Mito-BFP (red) and treated with 20μM CCCP treatment at time 0 during live-cell imaging. Imaging conducted at the medial cell section. Corresponds to movie 9. Scale: 10μm. Time in mins. **(B**) Quantification of colocalized Parkin signal with mitochondrial signal in live-cell imaging after 20μM treatment of CCCP at time 0 for control (black) and myc- VAP-B overexpressing (red) U2OS cells. N = 29 cells for ctrl and 28 for VAPB overexpression. 2 independent experiments. Error ± S.E.M. **(C)** Scatter plot of Parkin signal onset on mitochondria for control and myc-VAP-B overexpressing U2OS cells. Parkin onset time = 43.34 min ± 6.02 (mean ± s.d.) for control and 23.95 ± 8.578 for myc-VAP-B overexpression. Same dataset as in Figure 7B. P< 0.0001 (****). Student’s unpaired t-test. **(D)** Representative images of HeLa cells transiently expressing mitoBFP (red) and Parkin-GFP (green) either alone or in combination with ER-RFP-Mito (ER-Mito linker plasmid; cyan), treated with 20 µM CCCP at time point, fixed and imaged at various time points. **(E)** Quantification showing percentage of cells with Parkin positive mitochondria from images in 3D. N= 2 independent experiments having 50-100 cells from 15-27 fields for each condition. Error ± S.E.M. **(F)** Representative images of HeLa cells transiently expressing mitoBFP (red) and Parkin-GFP (green) either alone or in combination with ER-RFP-Mito (ER-Mito linker plasmid; lower panel) or PM-RFP-Mito (PM-Mito linker plasmid; middle panel) treated with or without 20 µM CCCP. **(G)**Quantification showing Parkin positive mitochondria area (%) in each cell from images in 3F. N= 2 independent experiments having Control (T0): 51 cells; ER-Mt (T0): 47 cells; PM-Mt (T0): 55 cells; Control (T45): 38 cells; ER-Mt (T45): 36 cells; PM-Mt (T45): 55 cells; Error ± S.D.

To corroborate the role of ERMCs in Parkin recruitment, we redirected mitochondria to the plasma membrane, by overexpressing a plasma-membrane mitochondrial linker construct (*PM- RFP-mito*) (Katona *et al*, 2022), that we predicted to decrease ERMCs. Accordingly, *PM-RFP-mito* reduces histamine-stimulated ER-to-mitochondrial calcium transfer (***Fig EV3C, 3D***). Also, *PM-RFP-mito* expressing cells lose the ability to accumulate Parkin on mitochondria, with 9.6 ±8.6% of their mitochondrial area positive for Parkin after 45-min treatment of FCCP (**Fig 3F, G**). Importantly, *PM-RFP-mito* does not inhibit CCCP induced ADA (***Fig EV3E***). Overall, these results suggest that (1) ER-mitochondrial proximity positively influences Parkin recruitment onto damaged mitochondria, and that (2) ADA delays Parkin recruitment by transiently disrupting ERMCs.

### ADA delays mitochondrial LC3 recruitment downstream of Parkin accumulation in an ERMC-dependent manner

Next, we asked whether ADA influences kinetics of LC3 recruitment downstream of Parkin accumulation. HeLa cells transfected with HA-Parkin display 9.0 ±5.0 % mitochondrial area positive for GFP-LC3 at 90 minutes after CCCP treatment (**Fig 4A, B**). CK666 pre-treated or Arp2 KD cells show significantly higher proportions of LC3-positive mitochondrial area at a similar time point (17.0 ± 5.9 % and 17.6 ± 5.4 % respectively) (**Fig 4A, B)**. Increasing ERMCs through overexpression of VAP-B or *ER-RFP-mito* phenocopies Arp2/3-inhibition, with 17.7 ± 4.5 % and 16.9 ± 5.3 % mitochondrial area positive for LC3 after 90-minute CCCP treatment respectively (**Fig. 4A, B**). As a control, we confirmed that CK666 pretreatment, Arp2 KD, VAP- B OE, or expression of *ER-RFP-mito* alone do not induce mitochondrial LC3 accumulation in the absence of CCCP stimulation (**Fig. 4A, B**). These results suggest that by delaying Parkin accumulation, ADA delays downstream mitophagy events, through regulation of ER-mitochondrial contacts.

**Figure 4:**
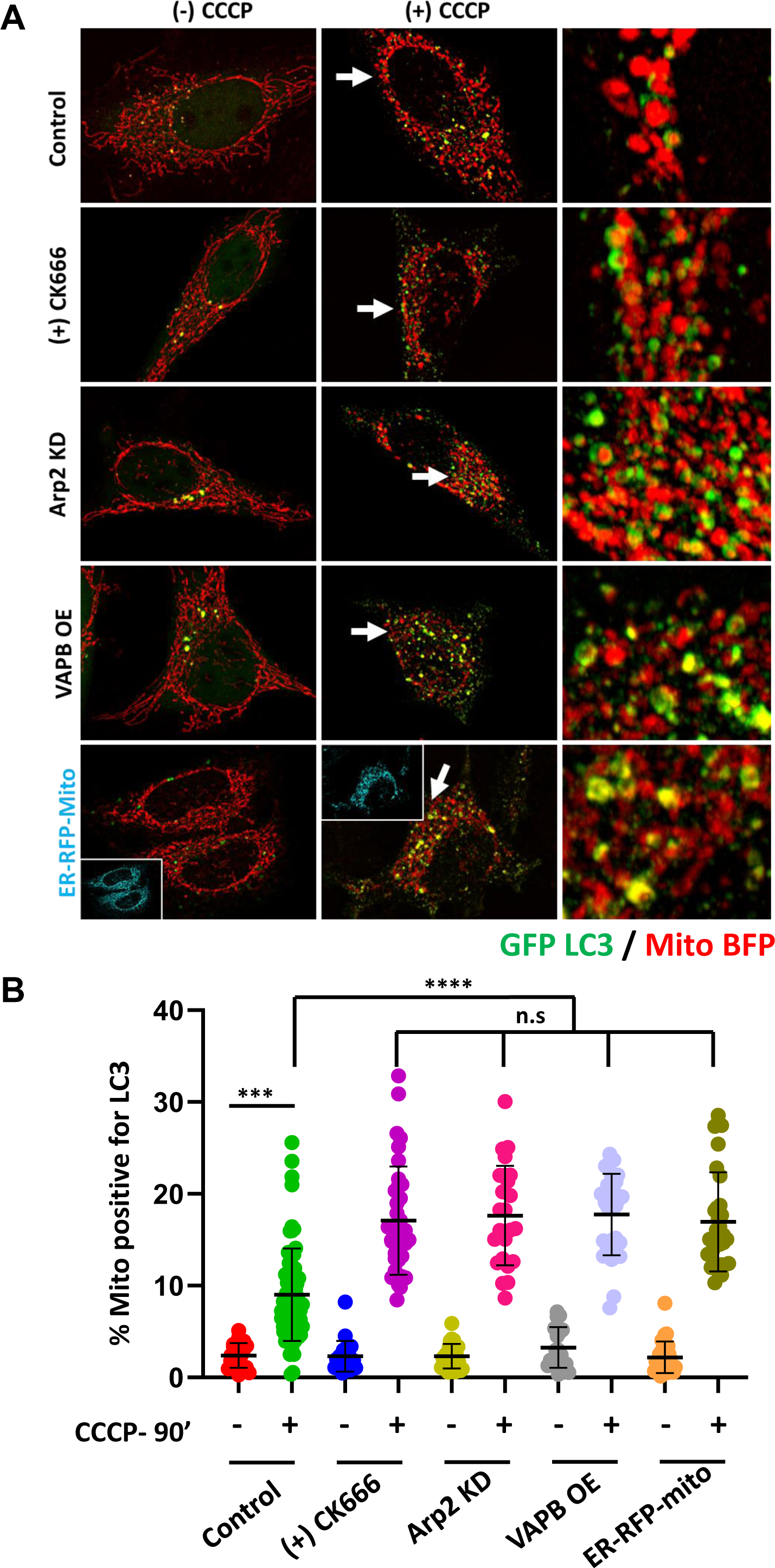
Enhancing ERMC speeds up LC3 recruitment onto depolarized mitochondria. **(A)** Representative images of control, CK666 treated (100 uM), Arp2 KD, VAP-B overexpressed and ER-RFP-Mito transfected HeLa cells transiently expressing GFP-LC3 (green), HA-Parkin and mitoBFP (red) before and after CCCP treatment (20 uM/ 90 mins). Regions next to arrows expanded (third panel) to show LC3 recruitment around mitochondria in CCCP treated cells from each condition. **(B)** Quantification showing LC3 positive mitochondria area (%) in each cell from images in 4A. N= 2 independent experiments having Control (untreated): 22 cells; CCCP 90’: 71 cells; CK666 90’: 21 cells; CK666 + CCCP 90’: 36 cells; Arp2 KD: 25 cells; Crp2 KD + CCCP 90’: 25 cells; VAP-B OE: 20 cells; VAP-B OE + CCCP 90’: 30 cells; ER-RFP-Mito: 28 cells; ER-RFP-Mito + CCCP 90’: 31 cells. P< 0.0001 (****) and P<0.001 (***) Error: ±SD

### Chronically dysfunctional mitochondria display actin-dependent mitophagy inhibition

We had previously shown that chronic OxPhos reduction through mtDNA depletion (achieved with a low dose of ethidium bromide (EtBr)) induces ADA-like peri-mitochondrial actin filaments (Chakrabarti *et al*, 2022). Despite persisting for days, these peri-mitochondrial actin filaments are eliminated by a 10-min treatment with CK666, showing that they are rapidly turning over and require continuous Arp2/3 complex activity (Chakrabarti *et al*, 2022). We therefore tested whether these ADA-like filaments inhibit both Parkin and LC3 recruitment onto chronically damaged mitochondria.

Treatment of MEFs with low doses of EtBr induces a significant reduction in mtDNA copy number (***Fig EV4A***), virtual elimination of both basal and stimulated OCR (***Fig EV4B***), and an increase in basal ECAR (glycolysis) (***Fig EV4B***). Similar to past results (Chakrabarti *et al*, 2022), mtDNA-depleted MEFs display peri-mitochondrial ADA-like filaments, sensitive to CK666 treatment (***Fig EV4C***). Next, we evaluated ER-mitochondrial proximity in these cells through PLA assay using antibodies against VAP-B (ER) and Tom20 (mitochondria). After 5 days of EtBr treatment we detected a 2-fold reduction in the PLA density compared to that of control cells (**Fig 5A, B**). This reduction is completely rescued by CK666 treatment (**Fig 5A, B**). These results suggest that ADA-like peri-mitochondrial actin filaments disrupt ERMCs downstream of mtDNA depletion mediated mitochondrial dysfunction.

**Figure 5:**
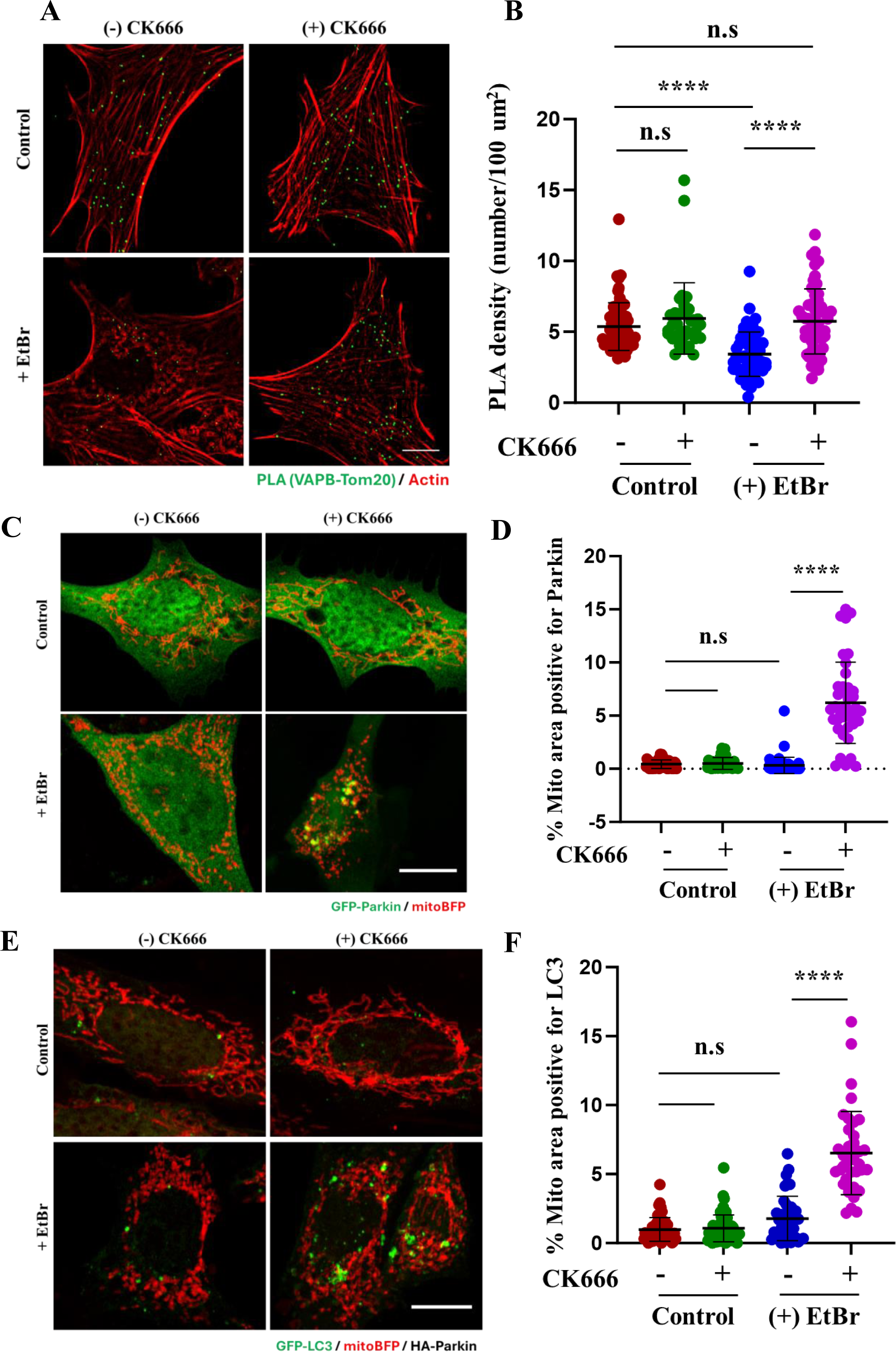
**Depletion of ADA like filaments in EtBr treated MEF cells induces ERMC and recruits Parkin and LC3 on chronically damaged mitochondria**. **(A)** Representative fixed-cell images from Proximity Ligation Assay (PLA; green) conducted between VAPB (ER) and Tom20 (mitochondria) to assess ERMC in control and EtBr treated (0.2 ug/ml for 5 days) HeLa cells treated with or without CK666 (100 μM/ 2 hr). Cells were stained with Phalloidin-488 to label actin filaments prior to PLA staining. Scale: 10 µm. **(B)** Scatter plot representing density of PLA dots (no. per 100µm^2^) in control and EtBr-treated HeLa cells with or without CK666 treatment. Same data set as Fig 5A. N= 65 cells (control); 35 cells (Control + CK666); 57 cells (EtBr Day 5) and 55 cells (EtBr Day 5 + CK666) from 20-25 fields and 2 independent experiments. P< 0.0001 (****); Error: ±SD **(C)** Representative fixed-cell images from control and EtBr treated (0.2 ug/ml for 5 days) HeLa cells transfected with Parkin-GFP (green) and mitoBFP (red), treated with or without CK666 (100 μM/ 4 hr). Scale: 10 µm. **(D)** Quantification showing Parkin positive mitochondria area (%) in each cell from images in 5C. N= 42 cells (control); 40 cells (Control + CK666); 58 cells (EtBr Day 5) and 40 cells (EtBr Day 5 + CK666) from 20-25 fields and 2 independent experiments. P< 0.0001 (****); Error:±SD **(E)** Representative fixed-cell images from control and EtBr treated (0.2 ug/ml for 5 days) HeLa cells transfected with LC3-GFP (green), mitoBFP (red) and HA-Parkin, treated with or without CK666 (100 μM/ 4 hr). Scale: 10 µm. **(F)** Quantification showing LC3 positive mitochondria area (%) in each cell from images in 5E. N= 54 cells (control); 61 cells (Control + CK666); 40 cells (EtBr Day 5) and 38 cells (EtBr Day 5 + CK666) from 20-25 fields and 2 independent experiments. P< 0.0001 (****); Error: ±SD

We also transfected these cells with GFP-Parkin or GFP-LC3 to evaluate their localization during chronic mitochondrial dysfunction. Surprisingly, EtBr-treated cells show little mitochondrial accumulation of both GFP-Parkin (**Fig 5C, D**) and GFP-LC3 (**Fig 5E, F**), in spite of these mitochondria being chronically depolarized. Interestingly, CK666 treatment causes a significant increase in both Parkin-positive (**Fig 5C, D)** and LC3-positive (**Fig 5E, F)** mitochondria in EtBr treated samples but not in the control samples. These results suggest that ADA-like filaments, assembled during chronic mitochondrial damage, disrupt ERMCs and block recruitment of Parkin and LC3 onto these damaged mitochondria. Removal of these ADA-like filaments restores ERMCs and allows Parkin and LC3 recruitment.

### Increasing the importance of mitochondrially-mediated ATP production causes prolonged ADA and delayed Parkin recruitment

The studies described above were conducted using standard culture media, which is hyperglycemic (25 mM glucose) compared to physiological glucose concentrations. Thus, cells cultured in this media – similar to cancer cells – mostly rely on aerobic glycolysis to meet their ATP demands (Bensinger & Christofk, 2012), make biosynthetic precursors(Shen *et al*, 2024; Bartman *et al*, 2023) and replenish NAD^+^(Luengo *et al*, 2021). Therefore, we wondered whether culturing cells under conditions requiring mitochondrial ATP production would affect ADA. We cultured HeLa cells for 10 days either in the presence of 10 mM glucose or in 10 mM galactose, the latter being known to cause a dependence on mitochondrial OxPhos for ATP production (MacVicar & Lane, 2014). Cells grown on glucose exhibit lower oligomycin-sensitive oxygen consumption rate (OCR) and higher basal extracellular acidification rate (ECAR) than galactose-grown cells (***Fig EV5A-C***). In addition, the maximal OCR for galactose-fed cells is higher than for those grown in glucose (***Fig EV5A-C***), suggesting that mitochondrial OxPhos is up-regulated in the absence of extracellular glucose. Mitochondrial polarization is similar in glucose- and galactose-primed HeLa cells (***Fig EV5D***), suggesting that mitochondria maintain appreciable Δψ_m_ in both cases.

Next, we compared ADA kinetics in the two conditions. Live-cell assays in U2OS cells show that CCCP-induced ADA is prolonged in galactose-primed cells compared to glucose-primed cells (**Fig 6A, *EV5E***). We confirmed this result with fixed-cell analysis by evaluating the number of HeLa cells displaying ADA at different time points following CCCP stimulation. We found a significantly higher proportion of ADA-positive cells at 5 and 10 min post-CCCP treatment in galactose-primed cells compared to glucose (**Fig 6B, *EV5F***).

**Figure 6:**
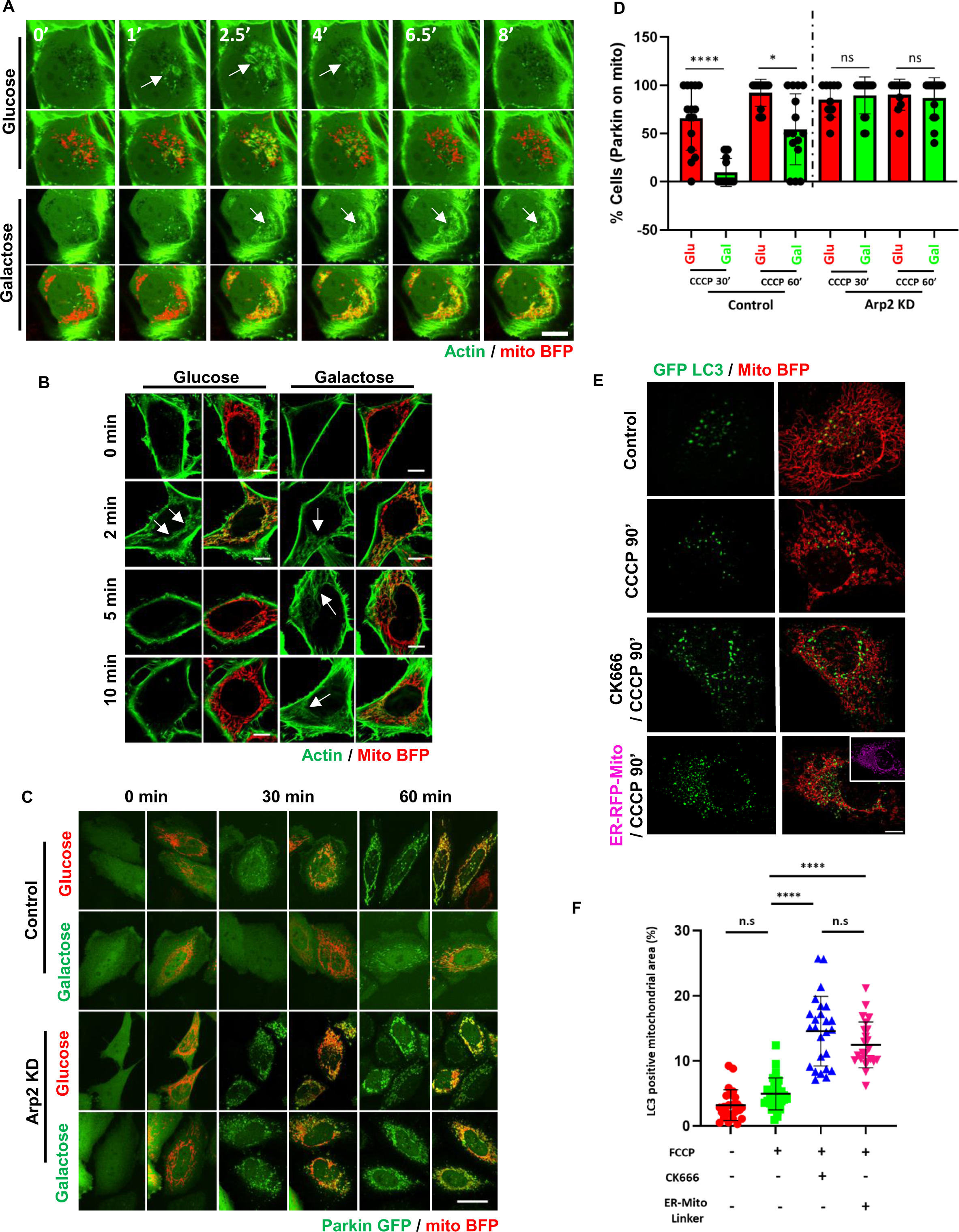
OxPhos-dependent cells display sustained ADA and inhibited Parkin recruitment. (A) Time-lapse montage of U2-OS cells cultured in 10 mM glucose or 10 mM galactose for 10 days transfected with GFP-Ftractin (actin; green) and mitoBFP (mitochondria, red) and treated with 20 uM CCCP at time 0. Imaging conducted at the medial cell section. Scale: 10μm. Time in mins. **(B)** Representative images of HeLa cells cultured in either 10 mM glucose or galactose for 10 days, transfected with mitoBFP (mitochondria, red), treated with 20 μM CCCP at time 0 and fixed at various time points. Cells were stained with rhodamine-phalloidin to show actin filaments (green) Scale: 10μm **(C)** Representative images of WT and Arp2 KD HeLa cells cultured in 10 mM glucose or galactose showing Parkin-GFP (green) and mitoBFP (mitochondria, red) treated with 20 μM CCCP and fixed at various time points. Scale: 10μm **(D)** Quantification showing percentage of cells with Parkin positive mitochondria from images in 6C. N= 2 independent experiments having 45-60 cells from 12-15 fields in each condition. Error ± S.D **(E)** Representative images of HeLa cells cultured in either 10 mM glucose or galactose for 10 days treated with FCCP (20 uM/ 90 min) with or without CK666 (100 uM) in the presence or absence of ER-RFP-mito (ER-mito linker). The cells were transiently transfected with GFP-LC3 (green), HA-Parkin and mitoBFP (red). **(F)** Quantification showing LC3 positive mitochondria area (%) in each cell from images in 6E. N= 2 independent experiments having Control (untreated): 22 cells; CCCP 90’: 32 cells; CK666 + CCCP 90’: 27 cells; ER-RFP-Mito + CCCP 90’: 26 cells. P< 0.0001 (****); Error: ±SD

We next compared CCCP-induced Parkin recruitment and downstream LC3 accumulation in glucose- and galactose-primed cells. Galactose-primed cells display reduced levels of mitochondrial GFP-Parkin following 30 min and 60 min of CCCP treatment, and this reduction is rescued by Arp2 KD. Similarly, mitochondrial levels of LC3 remain low after 90 minutes of CCCP in galactose primed cells (**Fig 6E, F**). Both Arp2/3 inhibition and overexpression of *ER-RFP-mito* significantly increased mitochondrial LC3 (**Fig 6E, F)**. Altogether, our data support a model whereby prolonged ADA induced by galactose-primed cells in turn resulst in a prolonged delay of Parkin-mediated mitophagy.

## Discussion

Collectively, the results presented in this study show that ADA has a significant effect on the kinetics of Parkin recruitment following mitochondrial damage. Acute mitochondrial depolarization generates a transient wave of ADA, which significantly delays the recruitment of Parkin onto depolarized mitochondria. Chronic mitochondrial dysfunction results in persistent ADA-like filaments that block Parkin recruitment, with actin depolymerization allowing rapid Parkin recruitment. Actin filaments do not merely form a barrier to prevent access but instead reduce ER-mitochondrial contacts. Overexpression of ER-mitochondrial tethers overcomes this ADA-induced block and accelerates depolarization-induced Parkin assembly in both the acute and chronic models of mitochondrial dysfunction. Our results also suggest that increasing the dependence of cells on OxPhos-derived ATP prolongs ADA, in turn causing prolonged suppression of Pink1/Parkin mediated mitophagy.

Actin filaments have diverse effects on ER-mitochondrial crosstalk. Previously we have shown that actin filaments polymerized on the ER, generated by the formin INF2, promote ERMCs, thereby stimulating ER-to-mitochondrial calcium transfer and mitochondrial division (Chakrabarti *et al*, 2018). Here, we show that a different form of actin filaments, polymerized by the Arp2/3 complex around dysfunctional mitochondria, acutely disrupts ERMCs. Actin filaments, therefore, play contrasting roles in regulating ERMCs, dependent on the cellular context.

The mechanism by which ADA disrupts ERMCs is unclear at present. Arp2/3-generated actin filament networks are known to generate force in many cellular contexts, such as at lamellipodia during cell migration (Krause & Gautreau, 2014) and in comet tails propelling intracellular pathogenic bacteria such as *Listeria* and *Shigella* (Rutenberg & Grant, 2001; Welch & Way, 2013). More recently, Arp2/3-mediated actin waves and comet tails around mitochondria during mitosis have been shown to regulate mitochondrial motility and distribution (Moore *et al*, 2021). A similar mechanism might be used by ADA to push ER and mitochondria apart. While the directionality of the growing actin filaments in ADA is still not clear, considering Arp2/3 recruitment to the mitochondrial membrane (Fung *et al*, 2022b), the force is likely to be directed toward the mitochondrion.

It is fascinating to note that distinct actin filament populations assemble around depolarized mitochondria at different times and for different purposes (Fung *et al*, 2023). While acute mitochondrial depolarization results in ADA within minutes, a second wave of actin occurs 1-2 hr post-depolarization, which we call PDA (Prolonged depolarization induced actin) (Kruppa *et al*, 2018). Both ADA and PDA are dependent on Arp2/3 but are distinct in the other proteins involved (WAVE Regulatory Complex and FMNL formins for ADA; N-WASP, myosin VI, and Parkin for PDA). Interestingly PDA appears to facilitate mitophagy in two ways: by preventing refusion of damaged mitochondria with healthy ones (Kruppa *et al*, 2018) and by dispersing clumps of damaged mitochondria (Hsieh & Yang, 2019). Finally, actin filaments are involved in several later steps of autophagy through a variety of Arp2/3 complex activators, including WHAMM (Kast *et al*, 2015; Kast & Dominguez, 2017), JMY (Hu & Mullins, 2019), and WASH (King *et al*, 2013). It is unclear whether PDA or autophagy-associated actin polymerization has effects on ERMCs.

The precise role of ERMCs in Parkin-mediated mitophagy is still unclear. Two promising candidates that might mediate this process are Pink1 and Mfn2, both of which accumulate at ERMC fractions following CCCP or Valinomycin treatment in multiple cell types, and both of which positively influence Parkin recruitment. Pink1 localization at ERMCs is required for BECN1 recruitment at the contact sites for later stages of mitophagy (Gelmetti *et al*, 2017). Moreover, the rapid stabilization of Pink1 (within 5 min of CCCP treatment) (Sekine *et al*, 2012) raises the possibility that ADA disruption of ERMCs might inhibit Pink1 stabilization. In addition to its critical role as an ER-Mitochondrial tethering factor (Naon *et al*, 2016; Chen *et al*, 2003; de Brito & Scorrano, 2008; Hu *et al*, 2021), Mfn2 is phosphorylated by Pink1 and functions as a receptor for Parkin (Chen & Dorn, 2013). In fact a progressive reduction of Mfn2, correlating with a reduction in mitophagy and accumulation of damaged mitochondria, has been linked to sarcopenia (Sebastián *et al*, 2016). However, the role of ERMCs in mitophagy might be more varied. The early stages of mitophagy might require an actual loosening of ERMCs to provide space for recruitment of the phagophore membranes, while some ERMCs might be retained as a source of phospholipid for the growing phagophore (Hamasaki *et al*, 2013; Gelmetti *et al*, 2017).

We show that cells cultured in galactose- containing media display prolonged ADA as well as a prolonged delay in Parkin recruitment upon depolarization, when compared to cells cultured in high glucose media. This result suggests that cells adapt to protect mitochondria once their ATP-producing function becomes essential. Moreover, we report that this delay in Parkin recruitment is eliminated by Arp2/3 inhibition, suggesting that the duration and persistence of peri-mitochondrial actin filaments regulates Parkin recruitment in this situation. Interestingly, studies have shown that OxPhos-dependent cells display impairment in OMA-1 dependent OPA-1 processing (MacVicar & Lane, 2014), depolarization-induced Parkin recruitment (Van Laar *et al*, 2011) and mitophagy (MacVicar & Lane, 2014). These impairments are reversed by culturing cells in glycolytic media conditions (Van Laar *et al*, 2011). Moreover, OxPhos-dependent cortical neurons do not accumulate Parkin upon acute mitochondrial damage (Van Laar *et al*, 2011). Coupled with our previous results showing that ADA inhibits OMA-1 dependent OPA-1 processing (Fung *et al*, 2022b, 2019), our present studies suggest that peri-mitochondrial actin assembly might be a key factor regulating early steps after acute mitochondrial dysfunction.

### Methods Cell culture

Wild-type human osteosarcoma U2OS and human cervical cancer HeLa cells were procured from American Type Culture Collection (ATCC). Mouse embryonic fibroblasts (MEF) were a gift from David Chan and is described elsewhere (Loson et al., 2013). All cells were grown in DMEM (Corning, 10-013-CV) supplemented with 10% newborn calf serum (Hyclone, SH30118.03) for U2OS or 10% fetal bovine serum (Sigma-Aldrich F4135) at 37°C with 5% CO2. Cell lines were tested every 3 months for mycoplasma contamination using Universal Mycoplasma detection kit (ATCC, 30-1012K) or MycoAlert Plus Mycoplasma Detection Ki (Lonza, LT07-701).

### DNA transfections, plasmids, and siRNA

For plasmid transfections, cells were seeded at 4×105 cells per well in a 35 mm dish at ∼16h before transfection. Transfections were performed in OPTI-MEM medium (Gibco, 31985062) using lipofectamine 2000 (Invitrogen, 11668) as per manufacturer’s protocol, followed by trypsinization and re-plating onto glass bottomed dishes (MatTek Corporation, P35G-1.5-14-C) at ∼1×105 cells per well (for live cell imaging) or coverslips (10 mm or 25 mm) for fixed cell analysis. Cells were imaged ∼16–24 h after transfection except for VAP-B overexpressing cells which were imaged within 8-12 hours after transfections.

GFP-F-tractin plasmid was a gift from C. Waterman and A. Pasapera (National Institutes of Health, Bethesda, MD) and were on a GFP-N1 backbone (Clonetech), as described previously (Johnson and Schell, 2009). mito-BFP (GFP-N1 backbone) constructs were previously described (Korobova et al., 2014) and consist of amino acids 1–22 of *S. cerevisiae* COX4 N terminal to the respective fusion protein. ERtagRFP (modified GFP-N1 backbone) was a gift from E. Snapp (Albert Einstein College of Medicine, New York, NY), with prolactin signal sequence at 5′ of the fluorescent protein and KDEL sequence at 3’. Myc-VAP-B was a gift from C.C.J. Miller (King’s College, London, UK) and described elsewhere (Gomez-Suaga et al., 2017). pLAMP1-mCherry was a gift from Amy Palmer (Addgene plasmid # 45147). cyto-CFP-FRBx5 was a gift from Takanari Inoue (Addgene plasmid # 103776). pEGFP-parkin WT was a gift from Edward Fon (Addgene plasmid # 45875). pMXs-IP HA-Parkin was a gift from Noboru Mizushima (Addgene plasmid # 38248). EGFP-LC3 was a gift from Karla Kirkegaard (Addgene plasmid # 11546). AKAP1-YFP-FKBP, ER-RFP-Mito, PM-RFP-Mito and mtGCampf6 constructs were a kind gift from Gyorgy Haznoczky (Thomas Jefferson University, Philadelphia, USA).

The following amounts of DNA were transfected per well (individually or combined for co transfection): 500 ng for mito–BFP, GFP–F-tractin, ER-RFP-Mito, PM-RFP, Mito, mtGCampf6 and pLAMP1-mCherry; 100 ng for pEGFP-parkin WT, pMXs-IP HA-Parkin and AKAP1-YFP-FKBP; 250 ng for GFP-LC3; 800 ng for ER-RFP; 750ng for Cyto-CFP-FRBX5; 400 ng for myc- VAP-B.

For all siRNA transfections, 1×105 cells were plated onto a 35mm dish and 2 ul RNAimax (Invitrogen, 13778) with 63pg siRNA were used per well. Cells were analyzed 72-96h post siRNA transfection. For Arp2 siRNA transfections, 1×105 cells were plated directly onto glass-bottomed dishes (MatTek Corporation, P35G-1.5-14-C) or coverslips and 2 ul RNAimax (Invitrogen, 13778) with 63pg siRNA were used per dish. Cells were analyzed 48h for Arp2. For live-cell imaging, plasmids containing fluorescent markers were transfected into siRNA-treated cells 18–24 h prior to imaging, as described above.

siRNAs:

human FMNL1(IDT,hs.Ri.FMNL1.13.5): 5’-

GTGGTACATTCGGTGGATCATGTTCTCCACCGAAT -3’).

FMNL2 (IDT, hs.Ri.FMNL2.13.1, 5’-CATGATGCAGTTTAGTAA-3’). FMNL3 (Ambion, s40551, 5- GCATCAAGGAGACATATGA-3’).

Arp2 (IDT, custom synthesized, HSC.RNAI.N001005386.12.6, 5’-

GGAUAUAAUAUUGAGCAAGAGCAGA-3’); and

negative control (IDT, #51-01-14-04, 5’-CGUUAAUCGCGUAUAAUACGCGUAU-3′).

### Immunofluorescence

Cells (untransfected or transfected) were plated onto coverslips 16 h prior to fixation and staining. Following respective treatments, Cells were then fixed with either prewarmed 4% PFA (20 min) or 1% glutaraldehyde (10 min) at room temperature, enabling optimal preservation of the actin cytoskeleton. Cells were washed with PBS and glutaraldehyde fixed cells were additionally washed with NaBH4 (1 mg/ml; 3 X 15 min each). Cells were then permeabilized with either 0.1% Triton X-100 for 1 min (PFA fixed) or 0.25% Triton X-100 for 10 min (glutaraldehyde fixed) and again washed with PBS three times. Prior to antibody staining, cells were blocked with 10% FBS in PBS for ∼30 min. Primary antibody was diluted in 1% FBS/PBS and coverslips were incubated on a drop of antibody solution on parafilm in a wet chamber for 1 Cells were then washed 6 times with 1X PBS and appropriate secondary antibody was mixed with phalloidin (AF-488 or Rhodamine phalloidin) in 1% FBS/PBS was added to the coverslips and incubated for 1 hour. Coverslips were again washed 6 times with 1X PBS and mounted on glass slides using ProLong Gold antifade mounting media (Invitrogen #P36930).

Primary antibodies: anti-Tom 20 (Abcam ab78547) used at 1: 400.

Secondary antibody: Alexa Fluor 488-coupled anti-rabbit (Invitrogen #A11008), Alexa Fluor 647-coupled anti-rabbit (Invitrogen #A21245)used at 1:500.

Rhodamine Phalloidin (Invitrogen R415) used at 1:500; Alexa-Fluor 488-phalloidin (Invitrogen A12379) used at 1:500

### Proximity Ligation Assay

Duolink™ PLA technology was used to study the ER-Mitochondrial interaction. Duolink™ In Situ Red Starter Kit Mouse/Rabbit (MilliporeSigma, cat# DUO92101) was used. In brief, cells were cultured on 10 mm coverslips for 16 hours before drug treatments. Following respective drug treatments cells were fixed with prewarmed 4% PFA for 20 min at RT and washed with PBS (X 3 times). Cells were then permeabilized using 0.1% Triton-X-100 for 1 min, washed with PBS and incubated with AF488-Phalloidin (diluted in 1% FBS/PBS) for 1 hour in dark.

Cells were then washed with PBS and incubated with Duolink blocking buffer for 1 hour and incubated with primary antibodies (diluted in Duolink antibody diluent) for 2 hours at RT.

Primary antibodies used: anti-VAPB (Proteintech 66191-1-Ig), anti-PTPIP51 (Proteintech 20641-1-AP) and anti-Tom20 (Abcam ab78547). Cells were then washed with 5% BSA (15 min X 3) and incubated with Duolink secondary antibodies for 1 hr at 37C. Cells were then washed with Duolink wash buffer A (5 min X 3) and incubated with Duolink ligase for 30 min at 37C. Cells were further washed with Duolink wash buffer A (2 min X 3) and incubated with Duolink polymerase for 90 min at 37 C. The cells were finally washed with Duolink wash buffer B (10 min X 3) and mounted on glass slides using prolong gold antifade reagent. Images were acquired within 24-48 hours.

### Microscopy

Microscopy of fixed samples was performed on LSM 880 equipped with 63×/1.4 NA plan Apochromat oil objective using the Airyscan detectors (Carl Zeiss Microscopy). The Airyscan uses a 32-channel array of GaAsP detectors configured as 0.2 airy units per channel to collect the data that is subsequently processed using Zen2 software. Cells were imaged with the 405-nm laser and 450/30 filter for BFP, 488 nm and 525/30 for GFP, and 561 nm and 595/25 for RFP. Images for Fig 2C, 2E, 3D, 3F, 4A, 5A, 5C, 5E, 6E, EV1H, EV3E, EV4C were acquired using this system and quantified using Image J.

Fixed cell images for figures 2A, 3A,B, D, E and 6B,C were acquired using Dragonfly 302 spinning disk confocal (Andor Technology, Inc.) on a Nikon Ti-E base and equipped with an iXon Ultra 888 EMC CD camera, a Zyla 4.2 Mpixel sCMOS camera, and a Tokai Hit stage-top incubator was used. A solid-state 405 smart diode 100-mW laser, solid-state 488 OPSL smart laser 50-mW laser, solid-state 560 OPSL smart laser 50-mW laser, and solid-state 637 OPSL smart laser 140-mW laser were used (objective: 60× 1.4 NA CFI Plan Apo; Nikon). Images were acquired using Fusion software (Andor Technology, Inc.). Images acquired using this system and quantified using Image J.

Live cell imaging was conducted in DMEM (Gibco, 21063-029) with 25 mM D-glucose, 4 mM L-glutamine and 25 mM Hepes, supplemented with 10% newborn calf serum, hence referred to as “live cell imaging media”. Cells (∼3.5x105) were plated onto MatTek dishes 16 hrs prior to imaging. Medium was preequilibrated at 37°C and 5% CO2 before use. Dishes were imaged using the Dragonfly 302 spinning disk confocal (Andor Technology) on a Nikon Ti-E base and equipped with an iXon Ultra 888 EMCCD camera, a Zyla 4.2 Mpixel sCMOS camera, and a Tokai Hit stage-top incubator set at 37°C. A solid-state 405 smart diode 100 mW laser, solid state 560 OPSL smart laser 50 mW laser, and solid state 637 OPSL smart laser 140 mW laser were used. Objectives used were the CFI Plan Apochromat Lambda 100X/1.45 NA oil (Nikon, MRD01905) for all drug treatment live-cell assays; and CFI Plan Apochromat 60X/1.4 NA oil (Nikon, MRD01605) to observe transient depolarization events or Parkin recruitment during live-cell imaging. Images were acquired using Fusion software (Andor Technology, version 2.0.0.15). Parkin recruitment was imaged at the medial section of the cell.

For Parkin recruitment assay with drug treatments, cells were pre-treated with 1ml of live-cell medium containing 100μM CK666 (Sigma-Aldrich, SML006) (from a 20mM stock in DMSO) for 30min before imaging. During imaging, cells were treated with 1ml live-cell medium containing 40μM CCCP at the start of the third frame (4min, time interval set at 2min). Imaging was continued at least 1.5 hours with cells in medium containing a final concentration of 20μM CCCP and 50μM CK666. Control cells were pretreated with an equal volume of DMSO (replacing CK666) and stimulated with 20μM CCCP during imaging. To visualize more cells in the field, the 60×1.4 NA objective was used. Parkin recruitment assay in KD cells was similar, except without the pre-treatment step, acquisition time interval was either 1.5 or 2min. For cyto- CFP-FRBX5 recruitment studies, cells were transfected with respective plasmids, pre-treated with 1 ml of live cell media, and imaged for 5 frames at 12 sec/frame, following which either DMSO (Invitrogen, D12345) or CCCP (final 40 μM) was added and imaged for another 100 sec following which 1 ml of rapamycin (Sigma 553210) (final 10 μM)-containing live cell media was added and imaged for another 150 sec.

## Image analysis and quantification

### Quantification from live-cell imaging

Unless otherwise stated, all image analysis was performed on ImageJ Fiji (version 1.51n, National Institutes of Health). Cells that shrunk during imaging or exhibited signs of phototoxicity such as blebbing or vacuolization were excluded from analysis (maximal amount 10% for any treatment).

<u>Parkin recruitment</u>. Parkin association with mitochondria was analyzed in Fiji using the Colocalization and Time series analyzer V3 ImageJ plugin. Firstly, GFP-Parkin (rolling ball 30.0) and mito-BFP images (rolling ball 20.0) were background-subtracted and converted into 8- bit files. Parkin-associated mitochondria were thresholded using the Colocalization ImageJ plugin with the following parameters: ratio, 25% (0–100%); threshold channel 1, 25 (0–255); threshold channel 2, 25 (0–255); and display value, 255 (0–255) for U2OS; and ratio, 50% (0–100%); threshold channel 1, 50 (0–255); threshold channel 2, 50 (0–255); and display value, 255 (0–255) for HeLa. ROIs were drawn for individual cells in the overlapped pixel stack and analyzed using Time series analyzer V3. The ROI was selected as the bulk region of the cell containing mitochondria using the mito-BFP signal. Mean colocalized signal of Parkin on mitochondria was plotted with respect to imaging duration (1.5 hr) at 2min or 1.5min intervals. To calculate the time of initial Parkin onset, the time of clear deviation from the pre-treatment intensity was manually read from the individual curve of each cell. For trendline analysis, data points from the linear portion of the averaged kinetic curve were extracted and a trendline was calculated using Excel. The time for the trendline to cross the point where y = 0 was determined as the average time for Parkin onset.

<u>ER-Mitochondria, lysosome-mitochondria and FRB-mitochondria overlap</u>. U2OS cells transiently transfected with the respective markers were imaged live by spinning disc confocal fluorescence microscopy at 15-s intervals in a single focal plane at the medial section 2-4 μm from the base. ROIs in the peri-nuclear region (ER-mitochondria, lysosome-mitochondria) or for the whole mitochondrial network (FRB-mitochondria) were background-subtracted using “rolling ball radius” plugin of Image J with a value of 20.0. The respective channels were further processed for bleach correction in Image J using “simple ratio” and converted to 8-bit images. The channels were then analyzed using the colocalization plugin in ImageJ with the following parameters: ratio 50% (0–100%); threshold channel 1: 30 (0–255); threshold channel 2: 1: 30 (0–255); display value: 255 (0–255) to obtain the overlapping pixels. The overlap intensity was then normalized to the pretreatment frames (1-5) and plotted over time.

### Quantification from fixed-cell imaging

<u>PLA assays</u>: All images were processed using Image J (NIH). z- projected images were background-subtracted using “rolling ball radius” plugin of Image J with a value of 20.0 and phalloidin (actin) channel was used to get the cellular area. Channel containing the PLA dots were converted to 8-bit and binarized. “Analyze particles” plugin was used to record total number of dots and presented as number of dots per 100 um2 of cellular area.

<u>Parkin and LC3 positive mitochondrial area:</u> z- projected images were background-subtracted using “rolling ball radius” plugin of Image J with a value of 20.0. The channels were then converted to 8-bit, Individual channels were thresholded using the Colocalization ImageJ plugin with the following parameters: ratio, 25% (0–100%); threshold channel 1, 25 (0–255); threshold channel 2, 25 (0–255); and display value, 255 (0–255) and overlapping pixel generated using the same plugin. Overlapping pixels were plotted as a function of total mitochondrial pixels.

### RT-qPCR

Total RNA was purified with NucleoSpin RNA Clean-up (Macherey-Nagel) following the manufacturer’s instructions. Genomic DNA was eliminated by on-column digestion with DNase I. 500 ng of RNA were reverse transcribed using LunaScript® RT Supermix (NEB) and qPCR (45 cycles) was performed on a QuantStudio 5 (Thermo Fisher Scientific). Reactions were run in triplicates with Luna® Universal qPCR Master Mix (NEB) in a total volume of 10 μl with standard cycling conditions. Relative gene expression was calculated using the ΔΔC_T_ method and normalized using ACTB as a housekeeping gene. All calculations were performed using QuantStudio Design and Analysis Software (Thermo Fisher Scientific). The list of primers used are as follows:

FMNL1 (F): 5’-CACCTGACCATCAAGCTGACC-3’ FMNL1 (R): 5’-CGTAGGCACATAATACAGACGTG-3’ FMNL2 (F): 5’-CAGGGAGCATGGATTCGCAG-3’ FMNL2 (R): 5’-TCAGGAGGTAGGTTCATAGCATT-3’ FMNL3 (F): 5’-TGGGGCTAGAGGAGTTCCTG-3’ FMNL3 (R): 5’-CCCCGACATCAAACACGTTG-3’ NCKAP1 (F): 5’-TTGTACCCCATAGCAAGTCTCT-3’ NCKAP1 (R): 5’-GGGCATTTCTCCACTGGTCAG-3’ Arp2 (F): 5’-CACCTGTGGGACTACACATTTG-3’ Arp2 (R): 5’-TGGTTGGGTTCATAGGAGGTTC-3’ ACTB (F): 5’-ACCTTCTACAATGAGCTGCG-3’ ACTB (R): 5’-CCTGGATAGCAACGTACATGG-3’

### Mitochondria DNA depletion and RT-PCR

mtDNA depletion was done as previously described (Chakrabarti, JCB 2022). Briefly, 1 × 10^5^ MEF cells were plated in a T-75 flask and incubated in DMEM + 10% FBS overnight. 24 h later, overnight media was replaced with either EtBr-containing media (DMEM+ 10% FBS + 0.2 μg/ml EtBr + 50 μg/ml uridine) or control media (DMEM + 10% FBS + 50 μg/ml uridine). Fresh media containing the chemicals were added every 48 hours. On the fifth day of treatment, cells were harvested to assess relative mtDNA copy number. Briefly, cells were trypsinized, resuspended in PBS buffer containing proteinase K (0.2 mg/ml), SDS (0.2%), EDTA (5 mM) and incubated for 6 h at 50 °C with constant shaking at 1,200 rpm. After isopropanol precipitation, DNA was resuspended in TE buffer and quantified by with a UV-Vis spectrophotometer (Implen). For the determination of mitochondrial DNA copy number, the genes encoding mitochondrial 12S and nuclear actin were amplified from 25 ng of DNA by qPCR on a QuantStudio 5 (Thermo Fisher Scientific) using standard cycling conditions (35 cycles). Reactions were run in triplicates using Luna® Universal qPCR Master Mix (NEB) in a total volume of 10 μl. Mitochondrial DNA copy number is defined as the relative amount of mitochondrial 12s normalized by a nuclear DNA normalization (ACTB), calculated with the ΔΔC_T_ method and QuantStudio Design and Analysis Software (Thermo Fisher Scientific). The list of primers used are as follows:

mt-12S (F): 5’-TAGCCCTAAACCTCAACAGT-3’

mt-12S (R): 5’-TGCGCTTACTTTGTAGCCTTCAT-3’ ACTB (F): 5’-TCACCCACACTGTGCCCATCTACGA-3’ ACTB (R): 5’-CAGCGGAACCGCTCATTGCCAATGG-3’

### Seahorse measurements

OCR and ECAR measurements were performed using the Seahorse XFe24 analyzer. In brief, 50,000 cells were seeded into each well excluding blank wells of 24-well cell culture miniplates suitable for the XFe24 analyzer. Concentrations of Oligomycin, FCCP, and Antimycin/Rotenone used at 1.5, 2, and 2.5/1 µM, respectively. Data for OCR, basal respiration, and ATP production in samples were calculated from Wave software (Agilent Technologies) and normalized to the corresponding total protein concentration.

### Lysates and Western blot

Cells from a 35mm dish were trypsinized, pelleted by centrifugation at 300 g for 5min and resuspended in 400μl of 1× DB (50mM Tris-HCl, pH 6.8, 2mM EDTA, 20% glycerol, 0.8% SDS, 0.02% Bromophenol Blue, 1000mM NaCl, 4M urea). Proteins were separated by SDS-PAGE in a Bio- Rad mini-gel system (7×8.4cm) and transferred onto polyvinylidene fluoride membrane (EMD Millipore, IPFL00010). The membrane was blocked with TBS-T (20 mM Tris-HCl, pH 7.6, 136 mM NaCl, 0.1% Tween32 20) containing 3% BSA (VWR Life Science, VWRV0332) for 1h, then incubated with primary antibody solution at 4°C overnight. After washing with TBS-T, the membrane was incubated with HRP conjugated secondary antibody for 1h at 23°C. Signals were detected by chemiluminescence. Primary antibody: anti-myc-tag antibody (Abcam, ab32 used 1:1000); anti-Myo IIA (CST #49349)

### Statistical analysis and graph plotting software

All statistical analyses and *P*-value determinations were conducted using GraphPad Prism QuickCalcs or GraphPad Prism 8 (version 8.3.0, GraphPad Software). To determine *P*-values, an unpaired Student’s *t*-test was performed between two groups of data, comparing full datasets as stated in the figure legends. For *P* values in multiple comparisons (unpaired), one-way Annova comparisons test was performed in GraphPad Prism 10. Live-cell actin burst and parkin curves, along with the standard errors of the mean (S.E.M) were plotted using Microsoft Excel for Office 365 (version 16.0.11231.20164, Microsoft Corporation).

## Acknowledgments

We thank David Chan for kindly providing the MEF cell lines, David Weaver for his expertise in imaging, Gyorgy Hajnoczky for his critical advice and reagents and Pak Rin for his ubiquitous presence behind the scenes. This work was supported by NIH R35 GM150811-1 (R.C), Margaret Q. Landenberger Research Foundation grant (R.C), Annesley Eye Brain Centre (R.C), NIH R35 GM122545-07 (H.N.H) and NIH R35147191-01 (M.T).

## Author Contribution

**Tak Shun Fung**: Data curation; Formal analysis; Investigation; Visualization; Methodology; Writing—review and editing. **Amrapali Ghosh**: Data curation; Formal analysis; Validation; Investigation. **Marco Tigano**: Data curation; Formal analysis; Validation; Investigation; Funding acquisition. **Henry Higgs:** Conceptualization; Resources; Supervision; Funding acquisition; Methodology; Writing—review and editing; Project administration. **Rajarshi Chakrabarti**: Conceptualization; Resources; Supervision; Funding acquisition; Methodology; Project administration; Writing—original draft; Writing—review and editing.

## Data availability section

This study includes no data deposited in external repositories

---

**Figure EV1:**
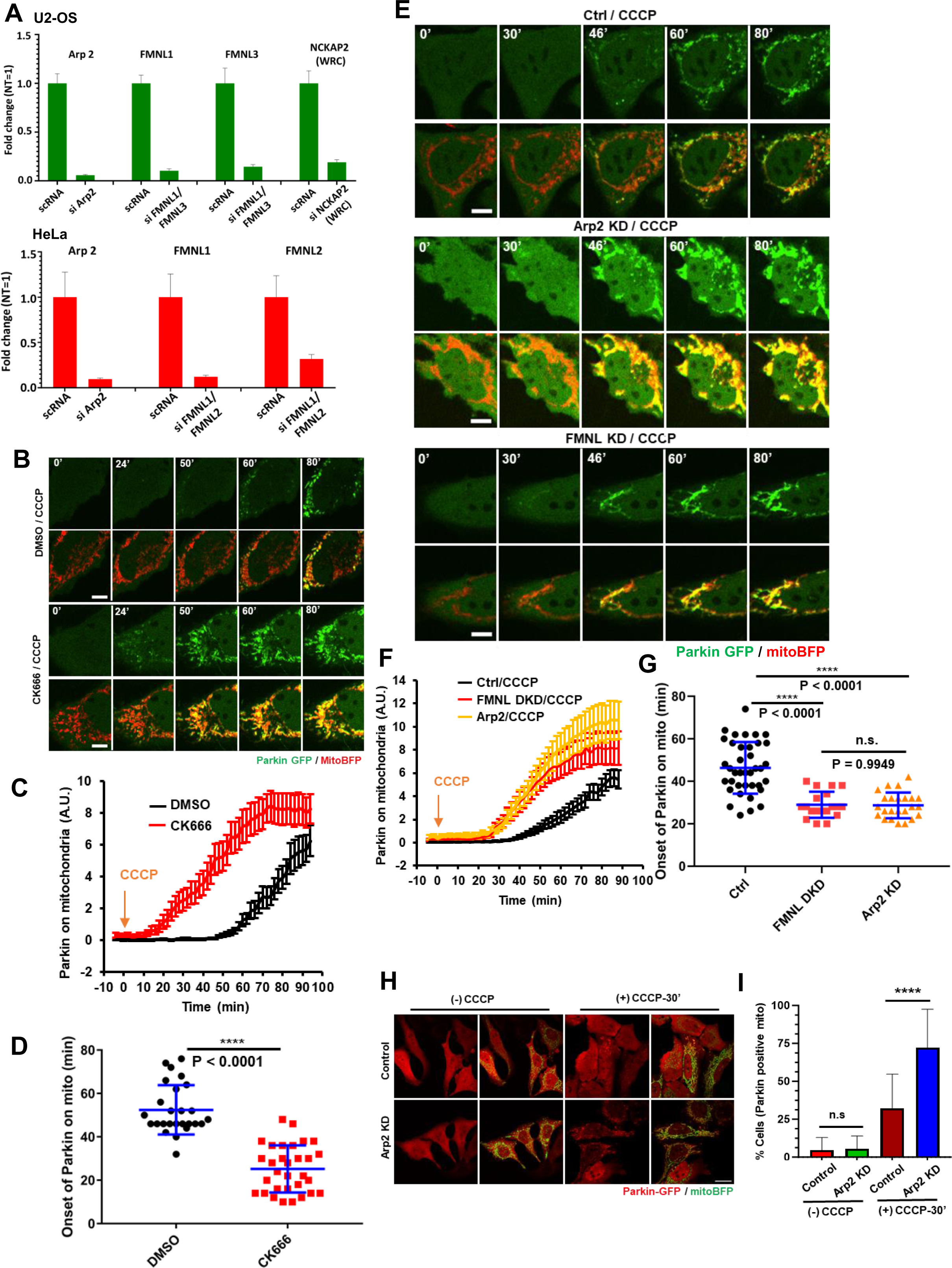
ADA delays Parkin recruitment on depolarized mitochondria. (A) RT-qPCR to validate knockdown of the respective genes in U2OS (upper; green bars) and HeLa cells (lower; red bars). Normalized expression data mean ± s.d. derived from *n* = 3 biological replicates. **(B)** Time lapse montages of WT HeLa cells transfected with GFP-Parkin (green) and Mito-BFP (red) pre-treated with 30 mins 100μM CK666 or DMSO before treatment with 20μM CCCP treatment during live-cell imaging at time 0. Imaging conducted at the medial cell section. Scale: 10μm. Time in mins. **(C)** Quantification of colocalized Parkin signal with mitochondrial signal in live-cell imaging after 20μM treatment of CCCP at time 0 for 30 mins DMSO (black) or 100μM CK666 (red) pre-treated HeLa cells. N = 26 cells for DMSO/CCCP and 31 for CK666/CCCP. 4 independent experiments. Error ± S.E.M. **(D)** Scatter plot of Parkin signal onset on mitochondria for DMSO or 100μM CK666 treated HeLa cells. Parkin onset time = 52.64 min ± 11.35 (mean ± s.d.) for DMSO/CCCP and 25.23 ± 10.89 for CK666/CCCP. Same dataset as in Figure S1C. P< 0.0001 (****). Student’s unpaired t-test. **(E)** Time lapse montages of ctrl, Arp2 KD, and FMNL1/2 DKD HeLa cells transfected with GFP-Parkin (green) and Mito-BFP (red) treated with 20μM CCCP treatment at time 0 during live-cell imaging. Imaging conducted at the medial cell section. Scale: 10μm. Time in mins. **(F)** Quantification of colocalized Parkin signal with mitochondrial signal in live-cell imaging after 20μM treatment of CCCP at time 0 for ctrl (black), FMNL1/2 DKD (red) or Arp2 KD (gold) HeLa cells. N = 38 cells for ctrl; 19 for FMNL1/2 DKD and 24 for Arp2 KD. 2 independent experiments. Error ± S.E.M. **(G)** Scatter plot of Parkin signal onset on mitochondria for ctrl, FMNL 1/2 DKD or Arp2 KD in HeLa cells. Parkin onset time = 46.32 min ± 12.16 (mean ± s.d.) for ctrl; 28.95 ± 6.12 for FMNL 1/2 DKD and 28.67 ± 6.01 for Arp2 KD cells. Same dataset as in Figure S1F. P< 0.0001 (****) for ctrl *vs* FMNL 1/2 DKD and ctrl *vs* Arp2 KD; P = 0.9949 (n.s.) for FMNL 1/2 DKD *vs* Arp2 KD. Tukey’s multiple comparisons test used. **(H)** Representative fixed cell images of WT and Arp2 KD HeLa cells transfected with Parkin-GFP (red) and mitoBFP (green) before and after 20 μM CCCP treatment. Scale: 10μm **(I)** Quantification showing percentage of cells with Parkin positive mitochondria from images in S1H. N= 2 independent experiments having Control: 278 cells/27 fields; Arp2 KD: 405 cells/35 fields; Control + CCCP: 191 cells/22 fields; Arp2 KD + CCCP: 141 cells/18 fields. Error ± S.E.M.

**Figure EV2:**
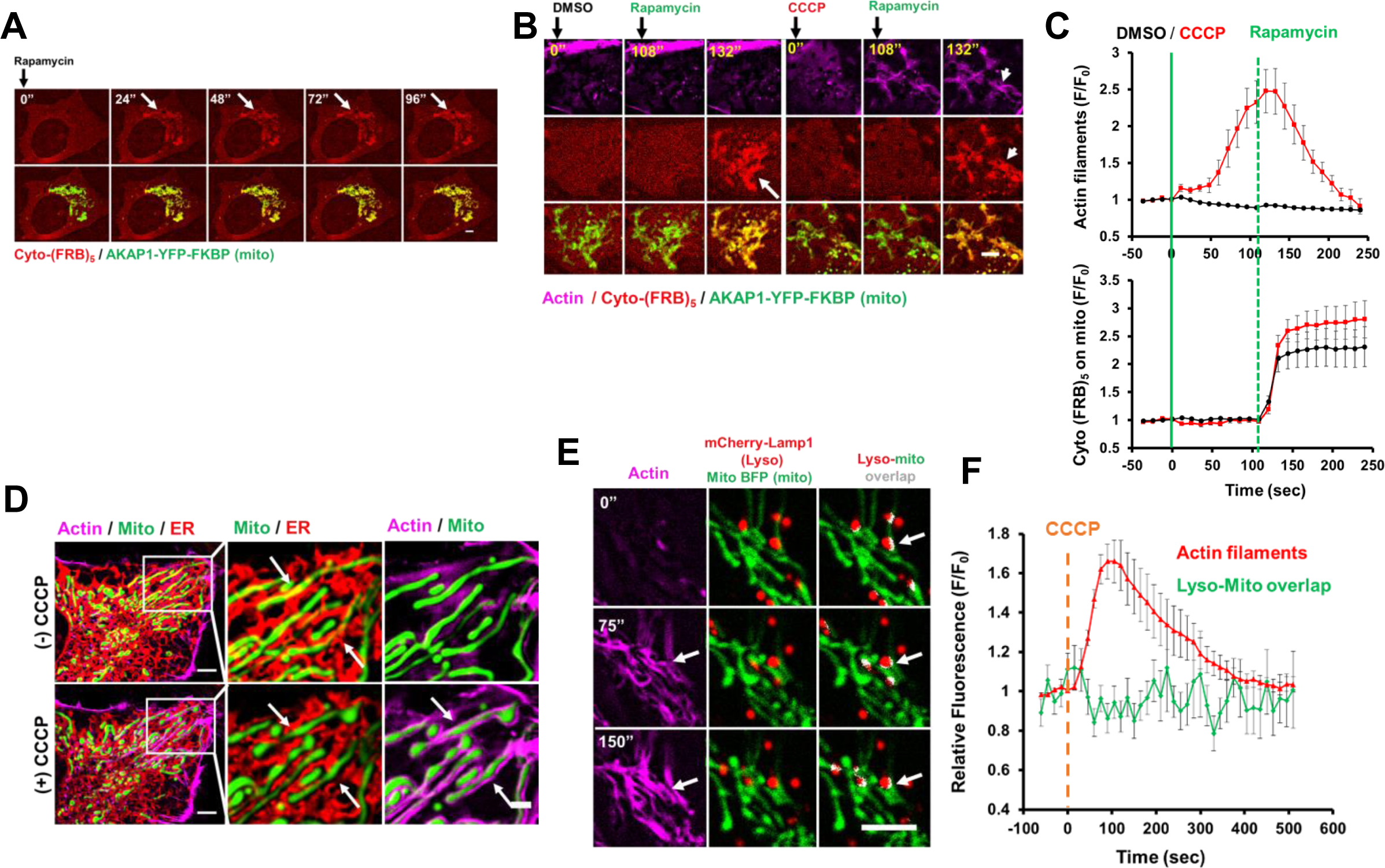
ADA disrupts ER-mitochondrial contacts but not mitochondria-lysosome contacts, (A) Time lapse montage of U2OS cells transfected with cyto-(FRB)5 and AKAP1-YFP-FKBP and treated with 10 μM Rapamycin at time 0 as indicated. Scale bar: 5 μm. **(B)** Time lapse montage of U2OS cells transfected with cyto-(FRB)5, AKAP1-YFP-FKBP CCCP and GFP-Ftractin and treated with either DMSO or CCCP for 100 seconds followed by rapamycin (final 10 μM). N= 9 cells from 2 independent experiments. **(C)** Quantification of actin filaments (upper) and mitochondrially associated CFP-(FRB)5 (lower) after the following treatments: DMSO or CCCP at time 0 followed by rapamycin (final 10 μM) treatment after 100 seconds of initial treatment. N= 9 cells from 2 independent experiments.**(D)** Micrographs from U2OS cells transfected with ERRFP (ER; red), mito-BFP (mitochondria; green) and GFP-F tractin (actin filaments; magenta) after 100 seconds of CCCP treatment. **(E)** Time-lapse montages of CCCP-induced actin polymerization and lysosomal-mitochondrial overlap in U2OS cells transfected with mCherry-Lamp1 (lysosomes; red), mito-BFP (mitochondria; green) and GFP-F tractin (actin filaments; magenta). **(F)**Quantification of CCCP-induced actin polymerization and Lysosome-mitochondrial overlap in U2OS cells transfected with mCherry-Lamp1 (lysosome; red), mito-BFP (mitochondria; green) and GFP-F tractin (actin filaments; magenta) N= 20 cells from 2 independent experiments.

**Figure EV3:**
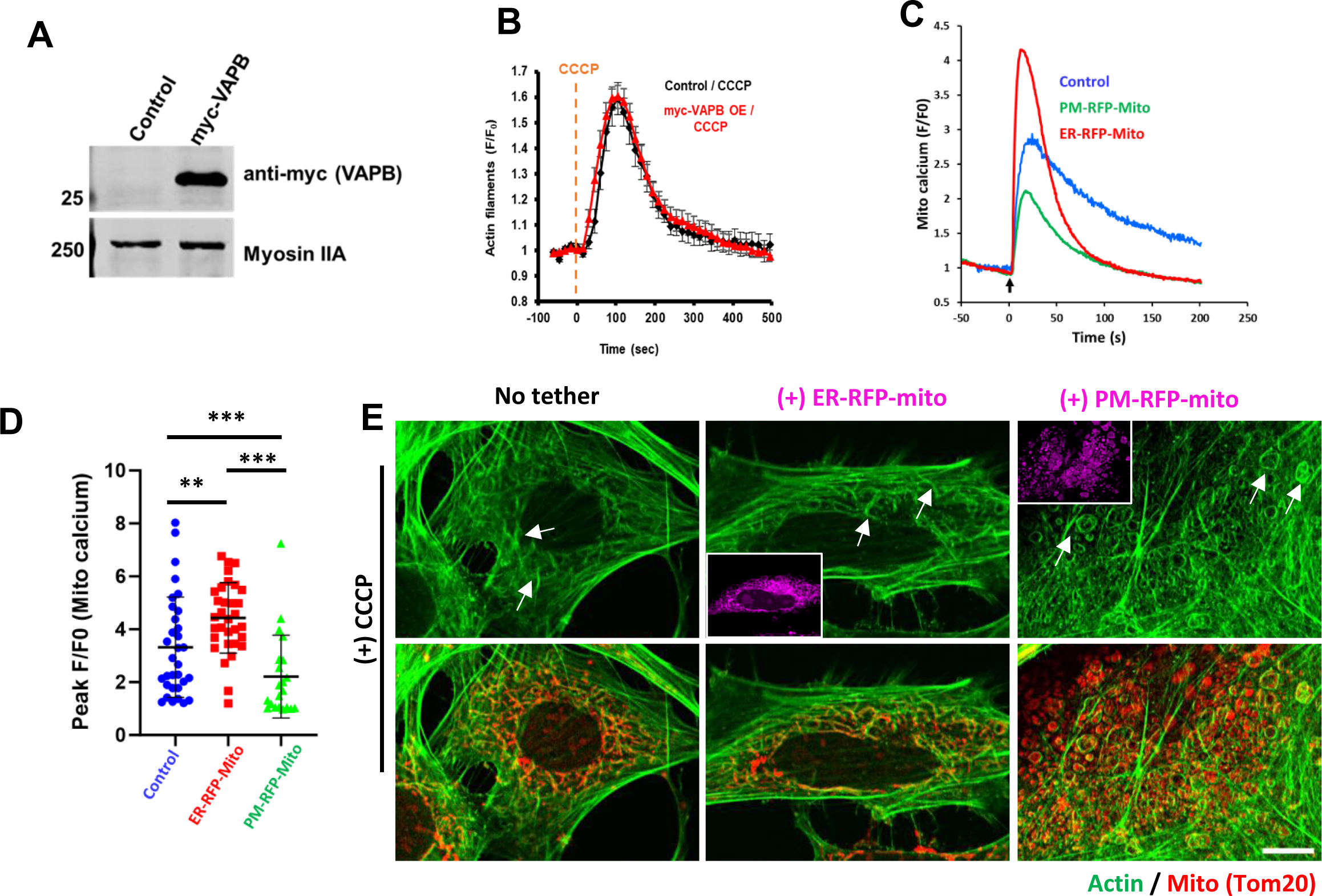
ERMC regulates CCCP induced Parkin recruitment, (A) Western blot for myc tag showing VAP-B expression in control and myc-VAP-B overexpressed U2OS cells. Myosin IIA is used as loading control. **(B)** Quantification of CCCP-induced actin polymerization in control and myc-VAP-B-overexpressing U2OS cells. N= 20 cells (control) and 45 cells (overexpressing myc-VAP-B). 2 independent experiments. Error ± S.E.M. **(C)** Averaged traces showing mitochondrial calcium fold change (MitoGCamp6f) following histamine stimulation (at time 0) in control (ER-RFP expressing), ER-RFP-mito (ER-Mito linker) and PM-RFP-Mito (PM-Mito-linker) transfected HeLa cells. Data from 2 independent experiments **(D)** Dot plot of peak fold change in mitochondrial calcium following histamine stimulation from individual traces as in Fig S3C. N= 32 cells (Control- ER-RFP); 32 cells (ER-RFP-Mito); 28 cells (PM-RFP-Mito). **(E)** Representative images of HeLa cells transiently expressing either ER-RFP-Mito (magenta) or PM-RFP-Mito, treated with 20 uM CCCP for 5 min, fixed and stained for actin (Rhodamine-phalloidin; green) and mitochondria (Tom20; red). Arrows show peri-mitochondrial actin assembly.

**Figure EV4:**
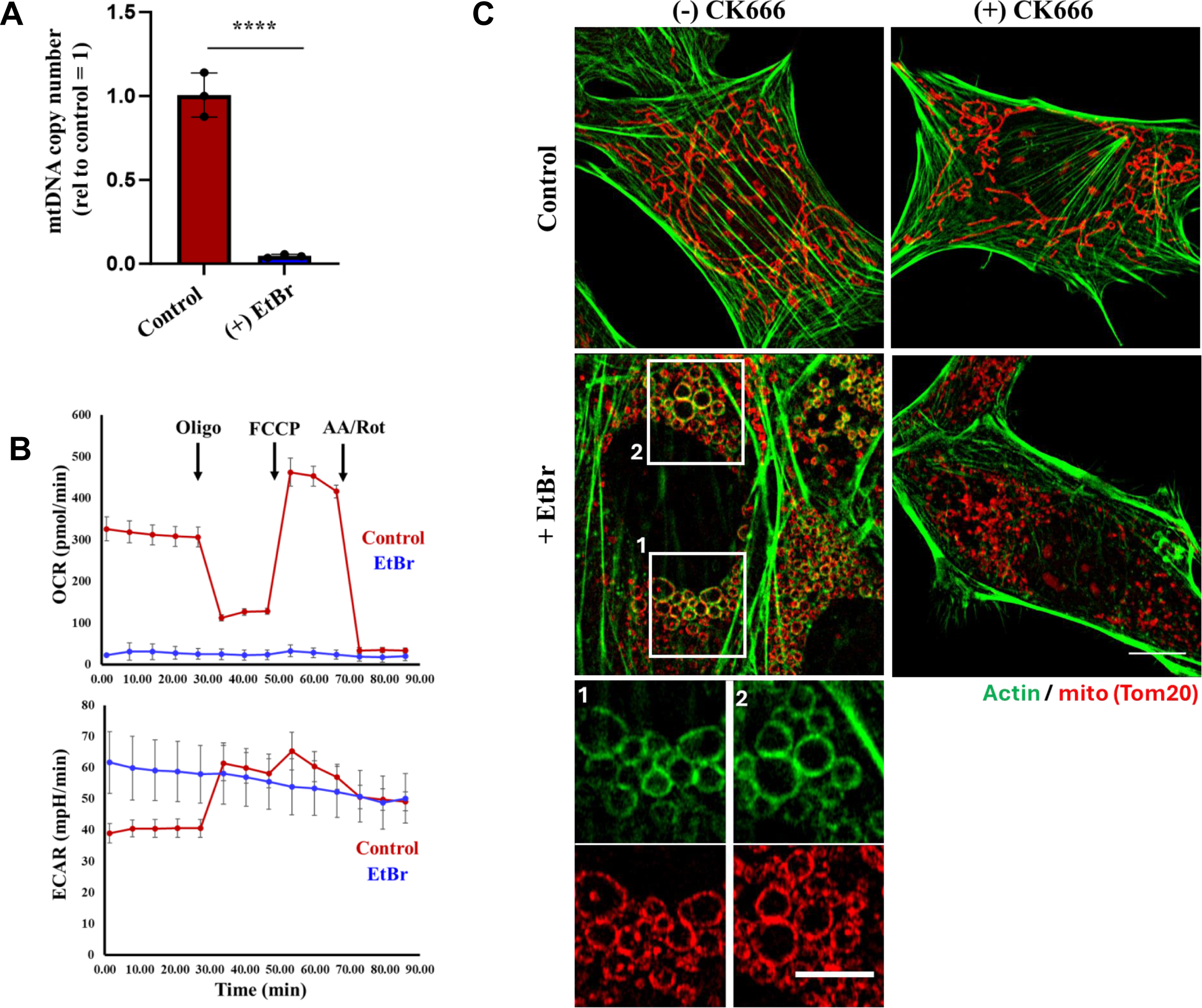
EtBr-induced mtDNA depletion and formation of ADA-like filaments. (A) RT-qPCR to validate depletion of mitochondrial DNA (mtDNA) in MEF cells treated with and without EtBr (0.2 ug/ml) for 5 days. Normalized expression data mean ± SD derived from *n* = 3 biological replicates. **(B)** OCR and ECAR traces from MEF cells treated with and without EtBr (0.2 ug/ml) for 5 days. Data from 2 biological replicates. **(C)** MEFs treated with or without EtBr (0.2 ug/ml) for 5 days, treated with CK666 (100 uM), fixed, and stained for actin filaments (green) and mitochondria (red). Scale bars: 10 μm.

**Figure EV5:**
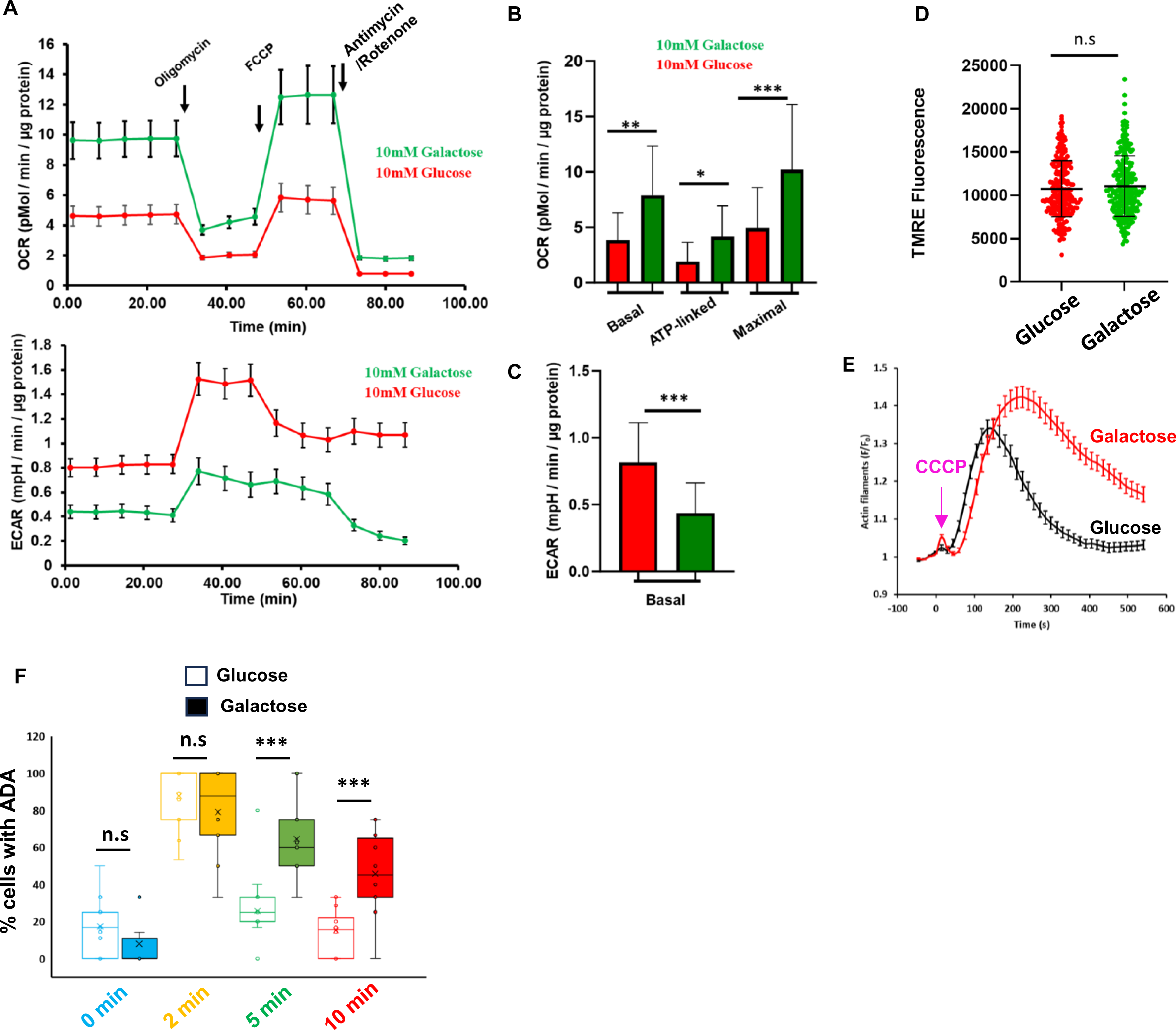
OxPhos-dependency causes prolonged ADA. (A) OCR and ECAR traces of HeLa cells cultured in either 10 mM glucose or galactose for 10 days, sequentially treated with oligomycin (1.5 uM), FCCP (1 uM) and Antimycin/Rotenone (2.5 uM/ 1 uM) at designated times. **(B)** Bar graph showing basal, ATP-linked, and maximal respiration calculated from traces shown in S5A. Error ± SD **(C)** Bar graph showing ECAR rates (glycolysis) in unstimulated HeLa cells cultured in glucose or galactose for 10 days from traces shown in S5A. Error ± SD **(D)** Dot plot showing TMRE fluorescence (measuring mitochondrial membrane potential) in HeLa cells cultured in either 10 mM glucose or galactose for 10 days. N= 213 cells (HeLa glucose) and 228 cells (HeLa galactose) from 2 independent experiments. Error ± SD **(E)** Graph of actin intensity (± SEM) around mitochondria in U2-OS cells cultured in glucose or galactose for 10 days as a function of time following 20 μM CCCP treatment. N= 97 cells (glucose); 69 cells (galactose) from 3 independent experiments. **(F)** Quantification showing percentage of cells with ADA in HeLa cells cultured in glucose or galactose (figures in 6B) following 20 uM CCCP treatment at time 0, fixed and stained for actin (rhodamine-phalloidin, green) and mitochondria (Tom 20, red) at various time points. N= 2 independent experiments having 40-80 cells from 12-15 fields in each condition. Error ± S.D

